# Heparinized Elastomeric Nanofibrillar Grafts: A Novel Approach for Mechanically Tunable, Cell-Supportive, and Thromboresistant Vascular Substitutes

**DOI:** 10.64898/2026.01.26.701857

**Authors:** Elizabeth C. Zermeno, John M. Kapitan, Alexander D. Sandquist, Amanda Reke, Apurbo Kumar Paul, Jason N. MacTaggart, Stephen A. Morin, Kaspars Maleckis

## Abstract

The clinical success of vascular grafts relies on three main prerequisites: artery-tuned mechanics, cell-supportive microstructure, and a thromboresistant interface. Most current solutions address only a subset of this triad and equate mechanical matching with compliance alone, which can lead to disturbed hemodynamics, maladaptive mechanobiology, and adverse graft-host biochemical interactions that frequently culminate in clinical complications and graft failure. This study presents polyurethane-based heparin-functionalized elastomeric nanofibrillar grafts (H-ENGs) that integrate all three prerequisites while allowing multi-parameter mechanical mimicry. To address the principal failure mode of early thrombosis, a small fraction of polyethyleneimine (PEI) is added to the ENG electrospinning solution to form P-ENGs, enabling one-step covalent heparin conjugation to form H-ENGs. The decoupled design of the ENG platform preserves the biomimetic microstructure and mechanics following PEI incorporation and heparinization, enabling adaptable, indication-specific optimization. In vitro, H-ENGs exhibit good cytocompatibility with minimal hemolysis, platelet adhesion, and whole blood clotting. Pilot porcine abdominal aorta interposition studies demonstrate feasibility: H-ENGs exhibit favorable surgical handling, intact suture-line integrity, and anastomotic hemostasis under dynamic flow, and retain artery-tuned mechanics and surface heparin at two weeks. While further testing is warranted, these results indicate that H-ENGs satisfy the three prerequisites for vascular graft clinical success.

**Graphical Abstract:** 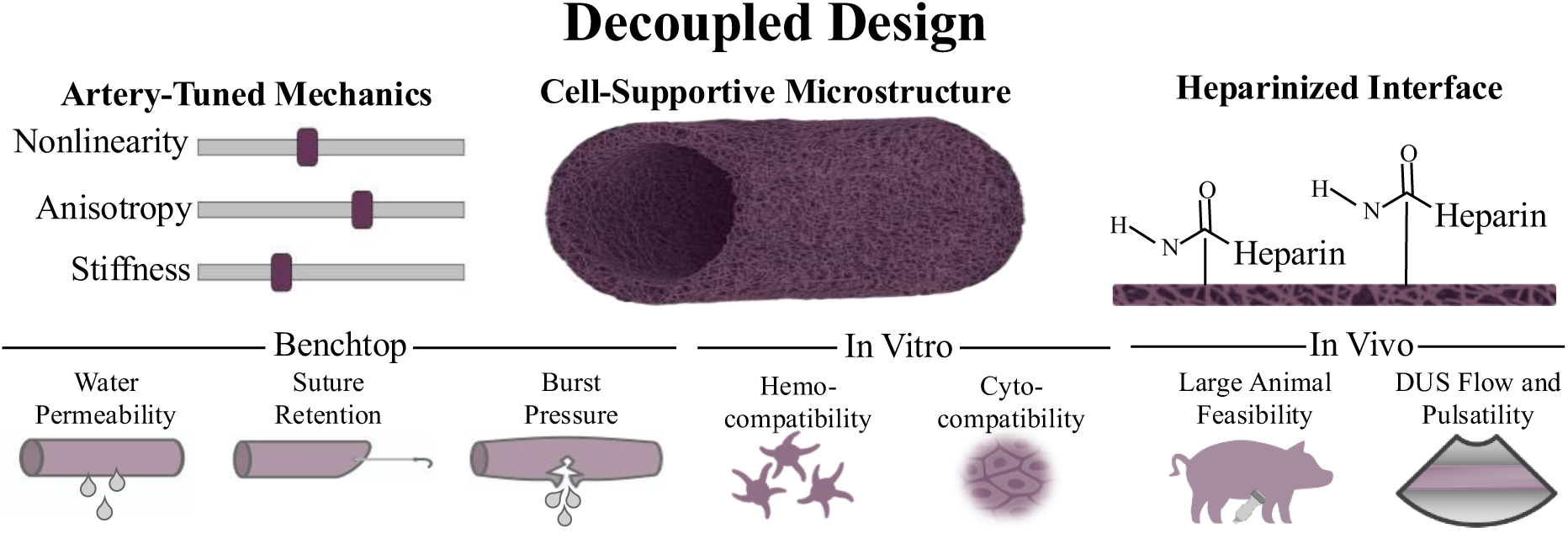

## 1. Introduction

Cardiovascular disease remains the leading cause of mortality worldwide, accounting for nearly 20 million deaths annually. *(1)* A substantial portion of this burden stems from atherosclerotic narrowing and occlusion of small- and medium-caliber arteries, which often necessitate revascularization to restore tissue and organ perfusion. For patients with advanced arterial obstructions, bypass grafting is often the only option to restore blood flow. Autologous vein grafts, typically the greater saphenous vein, remain the clinical gold standard due to its superior thromboresistance, cell-supportive microstructure, and biomimetic mechanical compliance. However, use of autologous substitutes is frequently limited by prior harvesting, comorbidities, and size constraints and often requires additional surgical procedures. *(2,3)* Even when available, vein grafts still lose patency at substantial rates within the first year. *(4,5)* These limitations underscore the enduring importance of effective synthetic vascular graft alternatives with off-the-shelf availability.

To offset the limited availability of autologous conduits, synthetic grafts made from expanded polytetrafluoroethylene (ePTFE), polyester (PET), and polyurethane (PU) are widely used, accounting for over half of the $1.7 billion vascular graft market. *(6–8)* Nevertheless, their long-term clinical performance remains substantially inferior to autologous vessels, with higher rates of complications and failure driven primarily by neointimal hyperplasia and thrombosis. *(2–5)* These shortcomings stem from the fundamental inability of existing synthetic materials to replicate the multifunctional properties of native arteries, resulting in chronically disturbed hemodynamics, limited functional regeneration of vascular tissue, and pathological host-graft biochemical processes. Addressing these limitations is non-trivial due to the complexity and specifics of arterial biomechanics, microstructure, and biochemical requirements. *(9–11)*

On the mechanical front, arteries exhibit nonlinearly elastomeric and anisotropic properties that vary substantially to support specific local anatomical and physiological requirements. *(6,12,13)* Elastomeric compliance supports large physiological deformations with high energy efficiency, while non-linear stiffening prevents overdistension and mechanical damage at elevated loads, and anisotropy accommodates the different mechanical requirements in circumferential and longitudinal directions. Large arteries favor low circumferential stiffness to store energy and buffer flow pulsatility, while smaller peripheral arteries require reduced axial stiffness and higher anisotropy to accommodate limb motion-induced deformations and minimize buckling. In contrast, most conventional synthetic grafts are stiff, linear, and isotropic, which prevents them from accommodating the aforementioned specific mechanical functions. *(2,4,5,14–16)* The mismatch in mechanical properties between the graft and the native artery has long been associated with increased rates of graft thrombosis and neointimal hyperplasia, driven by disturbed hemodynamics and mechanobiology that trigger endothelial dysfunction and smooth muscle cell over proliferation. *(2,3)* Despite longstanding efforts across the field to develop compliance-matched grafts, *(3,9,10,17)* compliance alone is an incomplete metric that neglects nonlinear stiffening and anisotropy of native arteries, thereby perpetuating mechanical mismatch. *(15,18,19)* This likely contributes to the limited clinical success of earlier compliance-matched grafts. *(20,21)* Moreover, even grafts with nonlinear, anisotropic elastomeric properties may still underperform if these parameters are not tuned to the specific artery. Elastomeric nanofibrillar grafts (ENGs), engineered through advanced electrospinning techniques to support multiparameter mechanical mimicry of arteries, were recently introduced, representing a significant advancement in mechanical optimization of synthetic arterial substitutes. These scaffolds demonstrated native artery-like deformations in vivo, thereby supporting improved vascular cell responses compared to commercial ePTFE controls. *(15)* Beyond mechanics, native arteries provide a complex and highly porous extracellular matrix structure that supports molecular transport, vascular cell migration and signaling, and adaptation to physiological changes. Similarly, microstructural architecture appears to strongly influence vascular tissue regeneration and function in synthetic grafts, which are essential prerequisites for long-term success. *(22–24)* Conventional ePTFE grafts, with limited pore interconnectivity, generally permit minimal tissue ingrowth, while PET grafts, though more porous, possess increased aneurysmal dilation and bleeding risks and still suffer from limited endothelialization in humans. *(25–35)* Electrospun scaffolds offer a compelling alternative, as advanced fabrication control now enables the tuning of fiber organization, pore size, and interconnectivity within the biologically optimal range for vascular cell migration and tissue integration, while minimizing excessive permeability. *(15,27,36–39)* Furthermore, the architectures of electrospun scaffolds also provide a more similar microenvironment to the extracellular matrix of vascular tissue and a larger surface area for vascular cell adhesion, *(40,41)* demonstrated in ENG vascular patch platform, which showed rapid smooth muscle cell infiltration and endothelialization within just two weeks. *(15)*

While mechanical and microstructural optimization play significant roles in optimal hemodynamics and vascular tissue function, these strategies alone cannot fully address one of the leading causes of graft failure: early thrombosis. *(42)* Most synthetic and natural graft materials remain inferior to native endothelium in terms of thromboresistance due to increased protein adsorption and platelet activation. To mitigate this risk of thrombosis before full coverage of native endothelium, a wide range of surface modification strategies has been investigated. Approaches such as heparinization, *(28,43–46)* nitric oxide–releasing coatings, *(47,48)* endothelial cell seeding, *(49,50)* and bioactive peptide modifications *(48,51)* have shown promise in vitro. However, in vivo success has been largely disappointing, as mechanical mismatch and suboptimal microstructure of these devices still cause hemodynamic disturbances and prevent the regeneration and viability of functional endothelium, which is the only established long-term defense against thrombosis. *(52,53)* This highlights the urgent need for novel integrated designs that address mechanical, microstructural, and biochemical prerequisites simultaneously. *(3)*

Here, we present a synthetic vascular graft platform that unifies multiparameter mechanical mimicry, cell-supportive microstructure, and enhanced thromboresistance. Unlike previous attempts that have largely focused on addressing individual failure causes at a time, our approach is a significant advancement in the field by providing a multi-prong solution to address all three main failure sources. The presented platform also combines the benefits of a streamlined manufacturing approach, design parameter decoupling, and tunability to a specific application. We demonstrate the efficacy of this approach through comprehensive mechanical, microstructural, and spectroscopic analyses, in vitro hemocompatibility and cytocompatibility assays, and pilot feasibility tests in a large animal model.

## 2. Results and Discussion

### 2.1. Graft Manufacturing

Building on our previous work, *(15)* the goal of this study was to employ a versatile route to add a stable antithrombogenic interface without constraining the ENG platform’s mechanical or microstructural design space. To that end, ENGs were manufactured according to Figure 1A, either with PEI (enabling subsequent heparin functionalization) or without PEI (non-functionalized controls). P-ENGs served only as an intermediate step in generating H-ENGs, as the presence of PEI provides primary amines required for covalent heparin attachment in later manufacturing steps (Figure 1B). Fourier transform infrared spectroscopy (FTIR) confirms the presence of these primary amine peaks at ∼3200 cm^-1^ (Figure S1). EDC/NHS coupling chemistry is then used to convert P-ENGs into H-ENGs (Figure 1B). *(54)* XPS analysis detects an S 2p signal characteristic of sulfated heparin on the surface of H-ENGs following functionalization and washing (Figure 1C), confirming successful stable surface immobilization of heparin. *(55)* The intent of this strategy is to covalently immobilize heparin on the graft surface, rather than to promote its release into circulation. This design follows established strategies in vascular grafts, where heparin is covalently immobilized to prevent rapid depletion and preserve antithrombogenic activity, *(56,57)* and ensures that bioactive heparin moieties remain stably presented at the blood-material interface, as confirmed by our XPS after extensive rinsing (Figure 1C). This simple process allows surface heparinization while maintaining the unique mechanical and microstructural properties of ENGs. While single-step heparinization is well established on planar polymer substrates such as ePTFE, *(58)* achieving stable immobilization on nanofibrillar networks has proven far more difficult. Their high surface area and ECM-like topology typically necessitate multistep processes (layer-by-layer assemblies, *(59)* polydopamine priming, *(60)* or plasma activation *(61)*) often involving coating precursors that are highly toxic and difficult to integrate. *(56)* Thus, this one-step heparinization method represents a promising advance for synthetic bypass grafts with ECM-like architecture.

**Figure 1.**
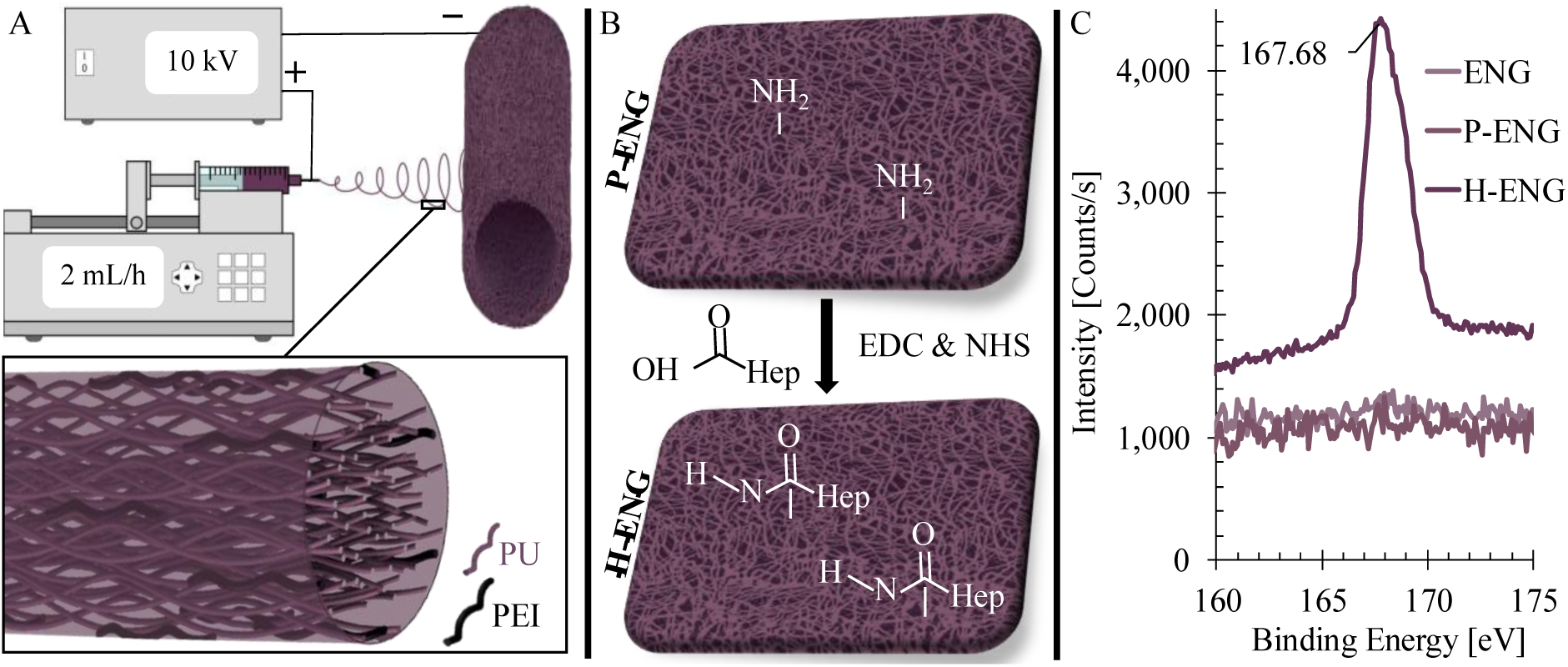
Heparinized Elastomeric Nanofibrillar Graft (H-ENG) manufacturing. A) Schematic of electrospinning PU solutions containing polyethylenimine (PEI) to produce PEI-integrated ENGs (P-ENGs). Inset: graphical illustration of PEI and PU molecules within the nanofiber. B) One-step heparin functionalization via EDC/NHS chemistry, in which primary amines from PEI enable covalent attachment of heparin carboxyl groups, yielding H-ENGs. C) S2p X-ray photoelectron spectroscopy (XPS) spectra for ENG, P-ENG, and H-ENG (n = 1 each), visualizing a strong S2p signal confirming successful heparin immobilization on H-ENGs.

### S2.2. Structure and Property Characterization of ENGs, P-ENGs, and H-ENGs

#### 2.2.1. Surgical, Microstructural, and Mechanical Evaluation

As a prerequisite for implantation, we established the surgical handling profile of the baseline ENG scaffold according to ISO 7198:2016 standard. Suture retention for both 45° and 90° cuts (2.84 ± 0.44 N and 2.91 ± 0.40 N, respectively) exceeds the widely accepted 2 N minimum threshold for cardiovascular grafts (Figure S2A). *(62)* Integral water permeability tests show an average of 45.73 ± 28.1 mL/min/cm^2^ at 120 mmHg. This indicates low graft permeability consistent with porous polymeric grafts designed for tissue integration and is well below the 800 mL/min/cm^2^ limit that would warrant pre-clotting of the graft. *(63,64)* Burst pressure strength testing was conducted utilizing a latex tubular liner. The latex liner failed after reaching pressures of 634 ± 66 mmHg, without rupturing the ENG, indicating ENG burst strength pressures exceed 634 mmHg (Figure S2B). This performance surpasses upper physiological limits of arterial pressures (typically < 200 mmHg), demonstrating sufficient mechanical integrity for vascular applications. *(65)* Together, these tests confirm that ENGs meet the basic surgical handling and safety requirements.

To ensure that our selected surface functionalization method preserves ENG mechanical and microstructural features, we quantified the morphology and mechanics of ENG variants both before and after functionalization. Bayesian analysis of SEM microstructure measurements and planar biaxial mechanical tests show no significant changes in fiber diameters, wall thickness, or axial and circumferential stiffness values after PEI incorporation and heparinization (Figure 2B, C, and Table S1). Collectively, the results demonstrate a substantive advance: a one-step heparinization that is decoupled from graft microstructure and mechanics, in contrast to earlier methods that report altered fiber diameter or stiffened scaffolds. *(66,67)*

**Figure 2.**
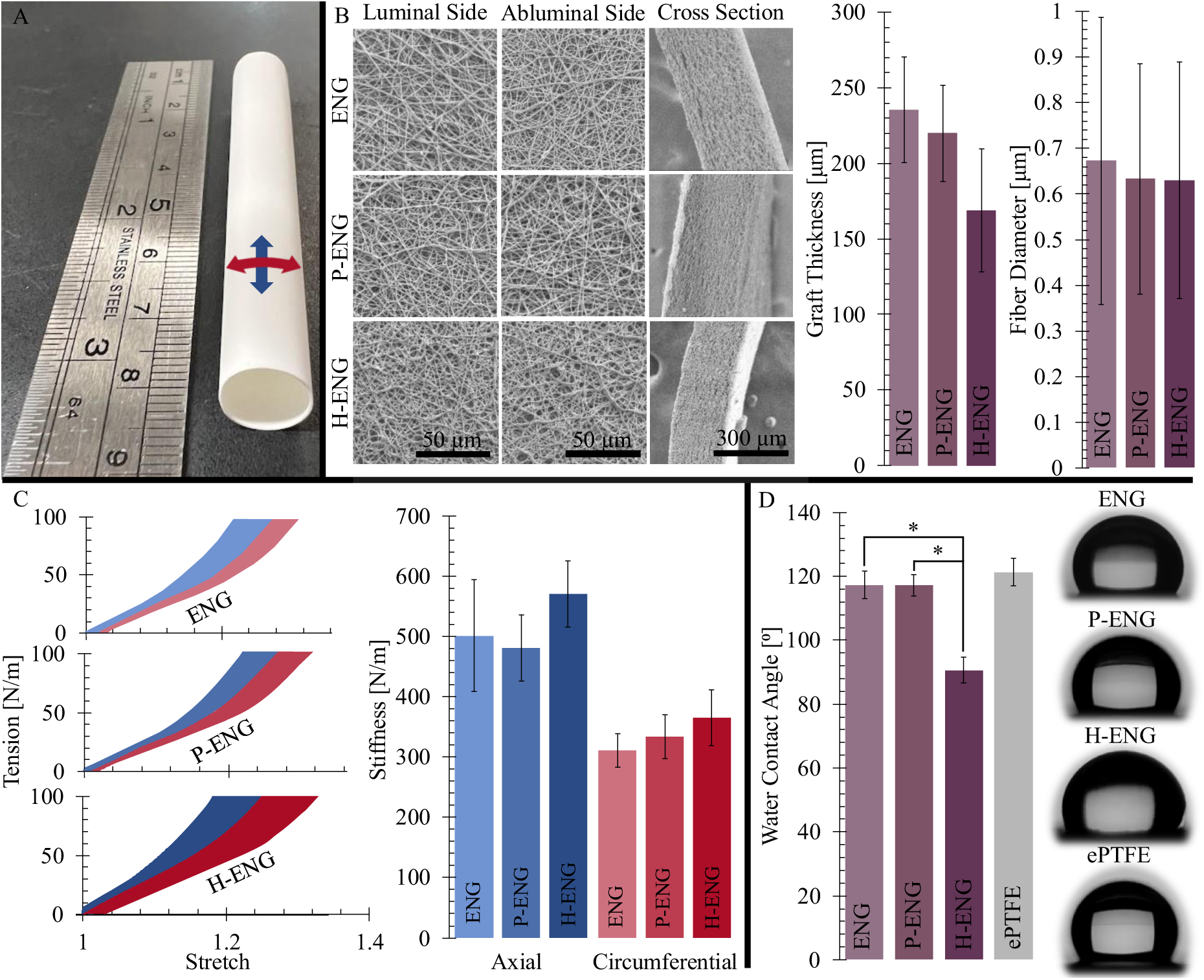
Geometric, microstructural, and physical characterization of ENGs, P-ENGs, and H-ENGs. A) Overall dimensions of the manufactured H-ENG device. B) Scanning electron microscopy (SEM) analysis of graft wall thickness and fiber diameters, with no significant differences among ENG variants. C) Planar biaxial mechanical response (left) and corresponding axial and circumferential stiffness values (right) of ENGs, P-ENGs, and H-ENGs show no significant differences. D) Static water contact angle measurements with representative images, showing significantly lower contact angle for H-ENGs compared to ENGs and P-ENGs. In all tests, n = 3 independent samples with three technical replicates each were characterized; error bars and shaded areas represent one standard deviation, and *p < 0.05.

#### 2.2.2. Water Contact Angle Measurements

Initial blood-graft interactions are governed by interfacial wettability and surface energy; thus, we first quantified water contact angle (WCA) and wetting kinetics as a rapid quality-control readout of heparinization and a coarse predictor of protein adsorption and initial cell adhesion on vascular grafts. *(68)* WCA measurements show that heparin functionalization profoundly increases surface wettability. Immediately after droplet deposition, WCA values were: 117.3 ± 4.3° for unmodified ENGs, 117.1 ± 3.4° for P-ENGs, 90.6 ± 4.1° for H-ENGs, and 121.3 ± 4.4° for ePTFE grafts. Bayesian regression detects no significant difference between ENGs and P-ENGs, indicating that the PEI linker alone does not alter the graft’s inherent, moderately hydrophobic character. H-ENGs, however, are significantly different from both ENGs and P-ENGs, exhibiting an almost 25% reduction in WCA (Figure 2D). While H-ENG’s immediate WCA lies outside reported optimal ranges, *(69)* the increased hydrophilicity of H-ENGs is still expected to favor endothelial and fibroblast attachment, *(70)* reduce nonspecific protein and platelet absorption, *(71)* and decrease thrombogenicity. *(69)*

Interestingly, the WCA of H-ENGs also exhibits time-dependent wetting kinetics. While ENG, P-ENG, and ePTFE angles remain essentially unchanged over four minutes, H-ENGs progress to complete wetting within this time frame (Figure S3). This behavior is attributable to the highly hydrophilic sulfate and carboxylate groups on surface-bound heparin, which promote water uptake and capillary imbibition into the fibrous network, *(72,73)* consistent with reports on other heparinized polymers. *(74)* Time-dependent increases in wettability have been linked to improved hemocompatibility *(75)* and endothelial adhesion, *(76)* suggesting a more optimal graft-blood interface once H-ENGs are fully wetted. Importantly, bypass grafts are routinely pre-soaked in heparinized saline before implantation, *(77)* so the fully wetted condition likely better reflects the in vivo interface suggesting low protein adsorption, limited platelet adhesion, and low thrombogenic potential for H-ENGs.

### 2.3. In Vitro Biological Compatibility Assessments

#### 2.3.1. In Vitro Hemocompatibility Assessments

While WCA is a useful first-pass screen for hemocompatibility, standards recommend assays with whole blood or components. The hemolysis assay (Figure 3A) shows that hemolysis rates in all samples are well below the 2% threshold, classifying them as non-hemolytic according to ASTM F756 standard. Platelet adhesion results (Figures 3A and S4) show that H-ENGs are not significantly different from ePTFE (40 ± 2 platelets/mm^2^ vs 44 ± 2 platelets/mm^2^, respectively). ePTFE is the most commonly used prosthetic material for peripheral bypass, *(78)* as such, it has been extensively tested in other literature and serves as a good benchmark with which to compare our materials. *(79,80)* Latex, assayed later as a pro-thrombotic positive control using the identical protocol, shows markedly higher platelet adhesion, although these data are displayed mostly for context. Complete descriptive statistics are available in Tables S2 and S3. P-ENG exhibits substantially and significantly higher platelet numbers compared to other material types, which is consistent with previous studies that show that amine groups from PEI incorporation increase platelet adhesion. *(81,82)* Surface heparin molecules in H-ENGs appear to successfully reverse this effect despite also containing PEI, presumably due to the anticoagulant nature of heparin and the reduced number of available amine groups as a result of EDC/NHS bonding. *(83)* Residual, unconjugated PEI remains a potential drawback of PEI-mediated heparin functionalization, although H-ENGs show no evidence of this effect in our assays. Results from the static whole-blood clotting assay appear to mirror those from the platelet-adhesion assays. After incubation, ePTFE and H-ENG show the lowest dry clot masses (0.1 ± 0.1 and 0.3 ± 0.8 mg/cm²), ENG shows low to moderate clotting (1.2 ± 0.4 mg/cm²), and P-ENG is markedly higher (7.0 ± 0.8 mg/cm²), consistent with protein adsorption-driven thrombosis. *(81–83)* Latex (PC) produces a large continuous clot (10.1 ± 0.9 mg/cm^2^), as expected for a pro-thrombotic positive control. *(84)* Together, these assays provide strong evidence for hemocompatible surface properties in H-ENGs.

**Figure 3.**
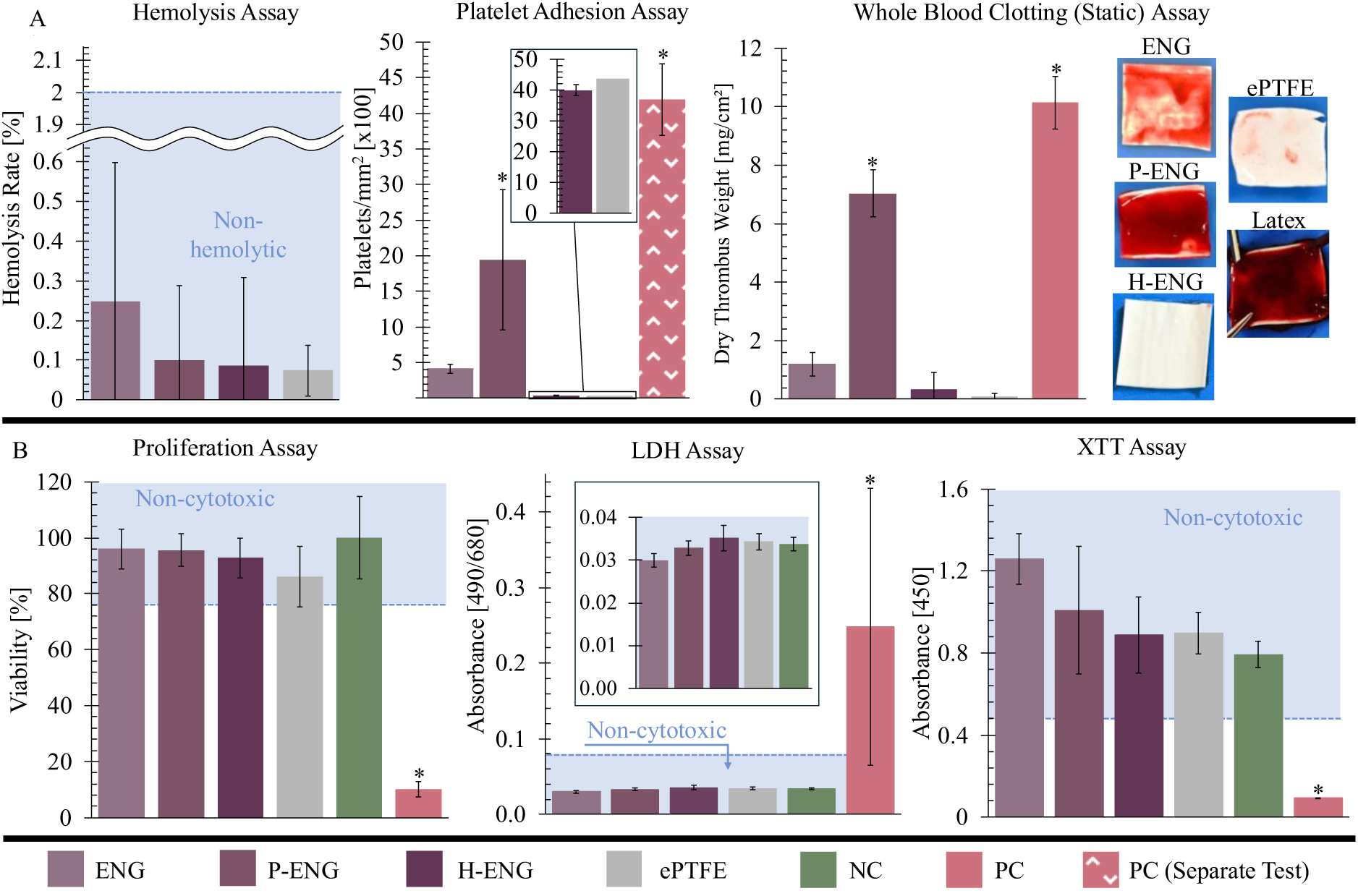
In vitro hemocompatibility and cytocompatibility assessments of ENGs, P-ENGs, and H-ENGs in comparison to commercial ePTFE. A) Hemolysis assay (left) indicates all grafts are non-hemolytic with no significant differences among groups; negative control (NC) = red blood cell (RBC) suspension, positive control (PC) = Triton X-100-treated RBCs. Platelet adhesion (middle) is low on ENGs and H-ENGs and comparable to ePTFE, whereas P-ENGs exhibit significantly higher adhesion. Platelet adhesion on latex (PC) was measured in a separate experiment using the identical assay conditions. Static whole-blood clotting assay (right) shows reduced clot formation on ENG and H-ENG relative to P-ENG, mirroring the platelet-adhesion trend. B) Cytocompatibility (HUVECs). Proliferation (left), LDH release (middle), and XTT metabolic activity (right) assays demonstrate non-cytotoxic performance across all groups; the LDH inset provides a magnified view of absorbance for all graft types and the assay NC. In all tests, n = 3 independent samples with three technical replicates each were characterized, error bars represent one standard deviation, and *p < 0.05.

#### 2.3.2. In Vitro Cytocompatibility Assessments

To further support translational safety, we assessed the cytocompatibility of ENG variants using HUVECs. Cell viability, as measured by the CyQUANT™ Cell Proliferation Assay, remains nearly 100% across all material samples (Figure 3B and Table S4). Similarly, the CyQUANT™ LDH Cytotoxicity Assay confirms the absence of cytotoxicity for all graft types, with values close to 0%. The XTT assay further supports these findings, showing high endothelial cell metabolic activity, though with slightly higher variability between samples. This variability may be attributed to the formazan-based assay’s sensitivity to pH changes in the growth medium, as cells are cultured in the same medium for approximately 36-48 hours. *(85)* Despite the increased variability, our statistical analysis supports the conclusion that all graft variants are non-cytotoxic. Previous studies report mild cytotoxic effects of PEI on HUVECs during extended culture. *(86)* In contrast, no such effects are observed in our experiments, likely due to the low PEI content (∼4.8% by weight) within the P-ENG and H-ENG fibers. As expected, ePTFE shows no cytotoxicity, consistent with its well-documented biocompatibility. *(87,88)* Immunofluorescence images further support favorable interactions between ENG variants and cells. HUVECs form confluent, homogeneous monolayers on all ENG variants (Figure S5). In contrast, cells on ePTFE grafts exhibit discontinuous growth patterns that follow the material’s surface topography, consistent with limited adhesion and migration on hydrophobic substrates. *(89)* 4’,6-Diamidino-2-phenylindole dihydrochloride (DAPI) staining confirms the presence of cell nuclei in ENG variants as well as ePTFE. However, the p120-catenin (p120) signal appears to be missing on the H-ENG. While this could suggest fewer mature adherens junctions at this time point, epitope masking or substrate-dependent signal interference are equally plausible explanations. *(90)* These short-term cultivation results demonstrate superior cell coverage and organization on ENG variants compared to commercial ePTFE control, suggesting that the nanofibrous architecture and surface chemistry of ENG variants support endothelialization, which is a critical factor for preventing late thrombosis in vivo.

### 2.4. In Vivo Feasibility Evaluation

Building on the in vitro hemocompatibility and cytocompatibility results, we performed an acute targeted pilot surgical feasibility evaluation in swine (n = 2). The objective of this feasibility study was not to estimate patency or efficacy but to evaluate surgical handling and stress-test early in vivo failure modes that are governed by material properties before neotissue forms, namely, suture-hole hemostasis, anastomotic suture-line integrity, and resistance to aneurysmal dilation in a high-flow vessel, as well as flow maintenance. We chose a two-week endpoint to capture interface-driven performance while endothelial coverage is minimal, thereby limiting confounding effects from tissue remodeling. This design yields an informative go/no-go decision while minimizing animal use and motivates a powered, longer-term study if feasibility is met.

At implantation, H-ENGs dilated appropriately without suture hole or perigraft bleeding (Figure 4A). At two weeks, both H-ENGs remained patent without occlusive thrombosis, infection, or aneurysmal degeneration. Ultrasound imaging shows a thin, flexible wall (Table S5 and S6) with pulsatility below 5% over the cardiac cycle (Figure 4B) over the two weeks. This modest pulsatility is likely due in part to sedation, which reduces cardiac output. *(91)* A single Bayesian regression across all four conditions (n = 2, 5 cycles per condition) confirms that there is no significant difference in pulsatility between the abdominal aorta at the time of implantation (2.29 ± 1.03%), the abdominal aorta at two weeks (1.72 ± 1.82%), the H-ENG (2.9 ± 0.74%) at the time of implantation, and the H-ENG at two weeks (1.99 ± 1.41%). These results confirm that H-ENGs maintain their artery-tuned, mechanical properties after two weeks in vivo.

**Figure 4.**
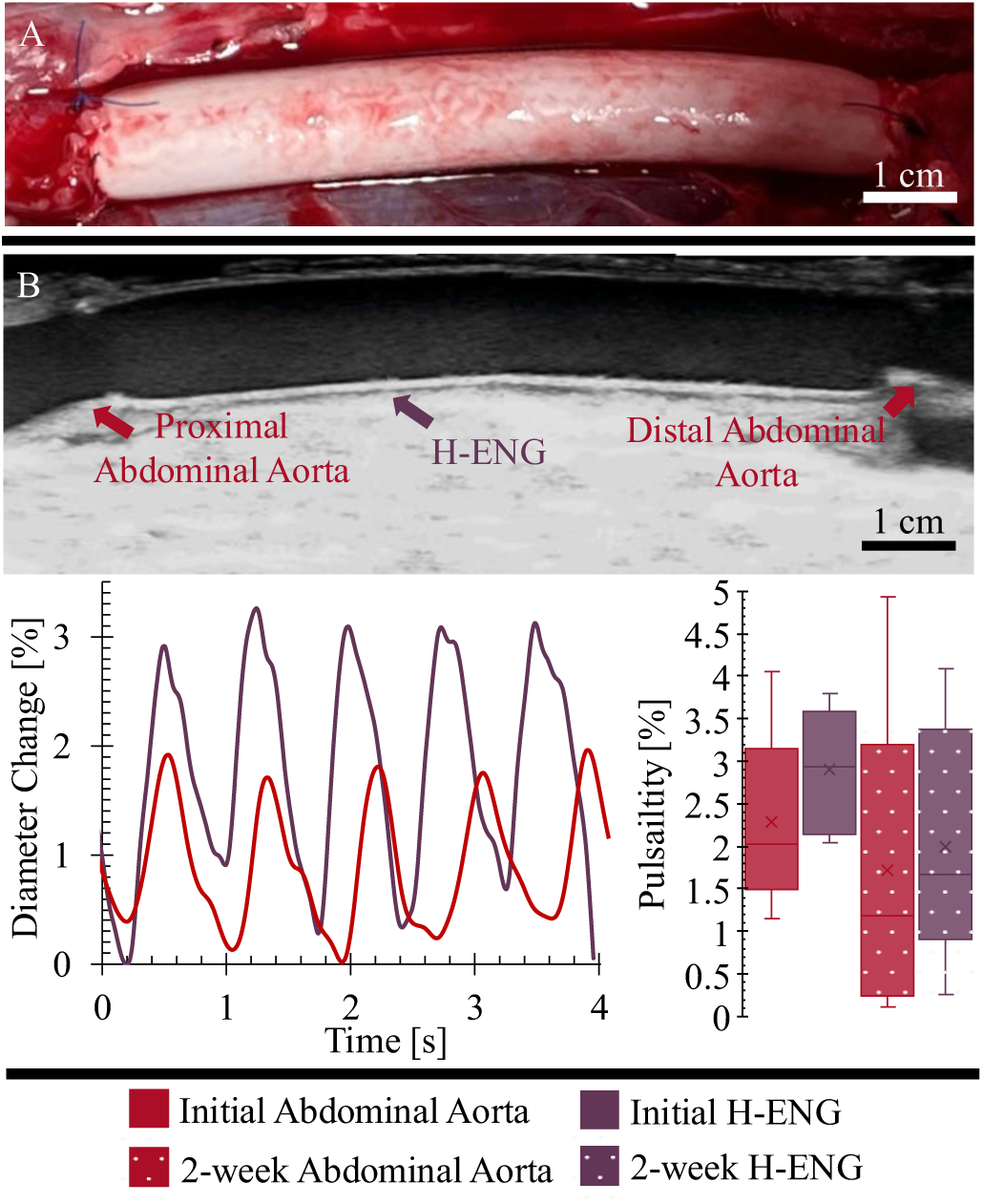
In vivo feasibility demonstration in the porcine abdominal aorta interposition model. A) Representative photo of H-ENG following implantation shows physiologic dilation with no evidence of suture line rupture or anastomotic bleeding. B) Representative longitudinal ultrasound still frame (top) with tracked relative diameter changes (bottom left) from five cardiac cycles of H-ENG and native abdominal aorta. Pulsatility measures (bottom right) at initial and two-week time points demonstrate comparable dynamic behavior between the native vessel and H-ENG. n = 2 independent samples with five technical replicates each, error bars represent one standard deviation, and *p < 0.05.

The observation that H-ENGs remain structurally intact and free from occlusive thrombi during this critical window provides strong evidence of short-term feasibility under high-pressure, high-flow arterial conditions. Importantly, unlike heparinized ePTFE grafts, which are notorious for excessive suture hole bleeding, *(92,93)* H-ENGs exhibit no such complications at implantation, underscoring their surgical manageability. Notably, while one localized non-occlusive thrombus was observed at an anastomotic site in one graft (visible near the distal anastomosis at the two-week ultrasound, Table S5), it did not propagate across the graft lumen, in contrast to the widespread thrombosis typically seen in less thromboresistant synthetic grafts *(94–96)*. Additionally, anastomotic thrombus is often not considered a reflection of graft surface properties but is rather attributed to changes in flow patterns. *(97)* The focal, non-propagating character is qualitatively consistent with our static whole-blood coagulation assay, in which H-ENGs showed reduced clot formation (Figure 3A). However, given the pilot scale and early timepoint, these observations are descriptive and do not establish in vivo antithrombogenicity or comparative performance.

### 2.5. Ex Vivo Evaluation

#### 2.5.1. Evaluation of Mechanical Properties Post-Implantation

To determine whether physiological exposure and early tissue ingrowth alter H-ENG mechanics, we performed planar biaxial testing on explanted grafts, as acute fibrin/cell infiltration can densify the fibrous network and bridge fibers, increasing effective stiffness and risking the reintroduction of mechanical mismatch. After two weeks, axial stiffness measures 469 ± 49 N/m versus 571 ± 55 N/m pre-implant, and circumferential stiffness 430 ± 15 N/m versus 365 ± 46 N/m pre-implant (Figure 5A). Statistical analysis indicates no significant differences between pre-implant and post-implant grafts, suggesting that biomimetic mechanical properties are maintained. Despite not being significant, the numerical trend of axial softening with circumferential stiffening may reflect early conditioning of the nanofibrous network by fibrin coating and initial cell ingress. Similar remodeling toward arterial-like nonlinearity and anisotropy was observed previously in non-heparinized ENGs. *(15)* These early trends motivate longer-term evaluation of in vivo mechanical evolution of H-ENGs.

**Figure 5.**
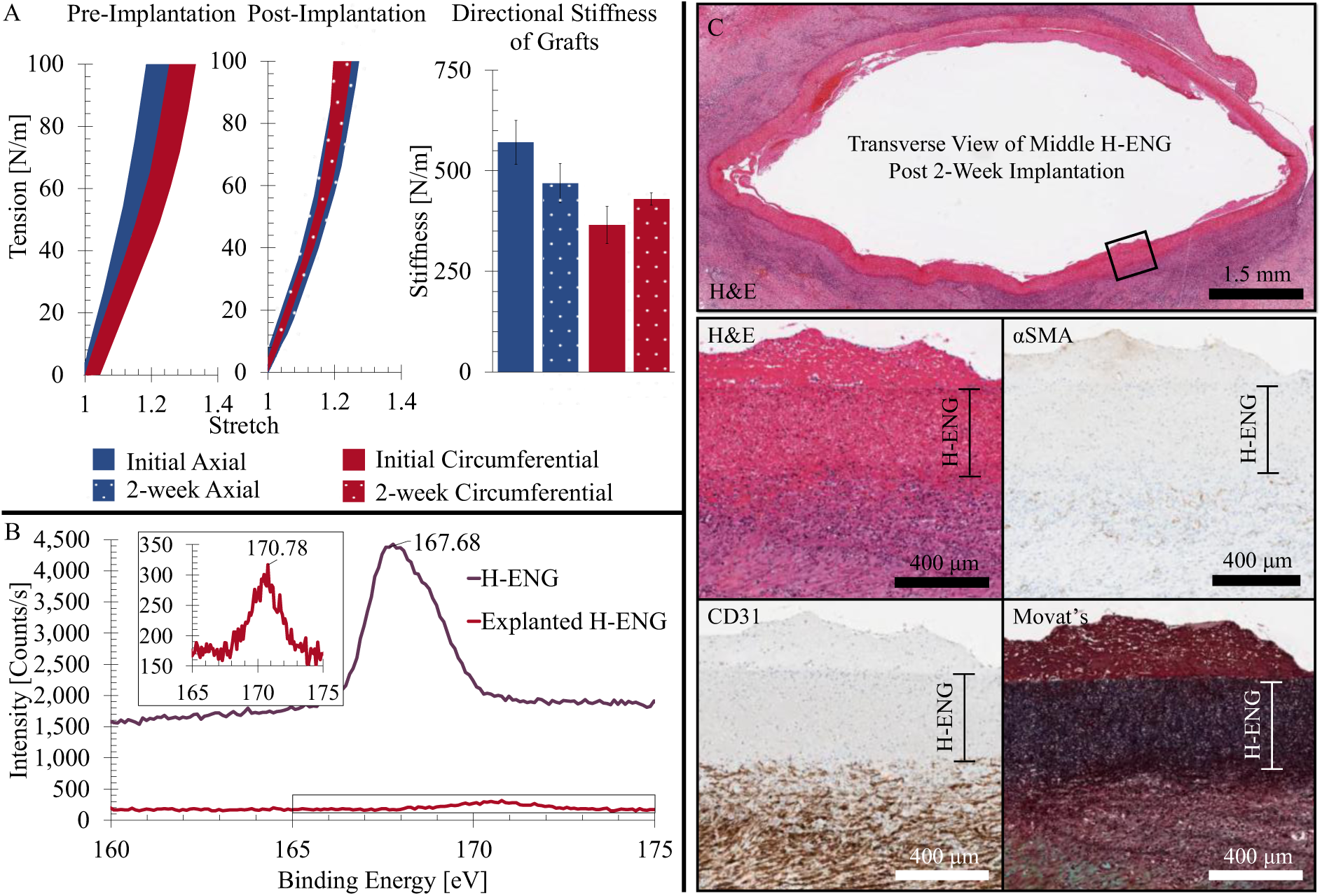
Ex vivo mechanical, chemical, and histological evaluation of H-ENGs. A) Planar equibiaxial stretch-tension response of H-ENGs before implantation (left, n = 3 with three technical replicates each) and post-implantation (middle, n = 2 with one technical replicate each). Shaded regions indicate the full range of intersample variability. Corresponding circumferential and axial stiffness values (right) indicate no significant changes pre-versus post-implantation. Error bars represent one standard deviation. B) Representative S2p XPS spectra of H-ENG pre- and post-implantation. The inset magnifies the S2p spectrum of the explanted H-ENG for better visualization of the sulfur signal. C) Representative histological images of transverse H-ENG stained with H&E, α-SMA, CD31, and Movat’s pentachrome stains (Movat’s), indicating minimal intimal thickening, limited smooth muscle cell ingrowth, and no meaningful luminal endothelialization.

#### 2.5.2. Verification of Surface Heparin Presence After Two Weeks in Vivo

Because antithrombogenic performance prior to full endothelialization depends on the stable presentation of heparin, we assessed surface chemistry after implantation. The XPS spectrum post-implantation exhibits a substantial decrease in the baseline and sulfur peak signal, which can be attributed to dried blood and adherent cells obscuring the underlying graft surface. The sulfur peak also shifts from 168.0 to 170.5 eV, consistent with changes in the binding environment due to the presence of copper or calcium ions commonly found in blood (Figure 5B). *(98,99)* Importantly, the characteristic S2p signal remains detectable after H-ENG explantation, confirming that covalently bound heparin was preserved throughout the two-week implantation period. This stability is achieved in an abdominal aortic interposition model that imposes high flow and pressure, with total flows substantially exceeding those in peripheral arteries. The persistence of immobilized heparin under these conditions supports the durability of PEI-mediated covalent functionalization in physiologic circulation and suggests that surface heparin in H-ENGs might be retained even longer at lower-flow arterial beds. Together, these findings provide evidence that the PEI-mediated covalent functionalization strategy confers durable antithrombogenic surface properties under physiologic circulation.

#### 2.5.3. Early Remodeling Evaluation Through Histological Imaging

To contextualize the mechanical and XPS findings, we examined early healing and remodeling by gross histological analysis. H-ENGs exhibit minimal intimal thickening, a response typically increased in mechanically mismatched grafts. α-SMA staining indicates limited abluminal smooth muscle cell ingrowth, while CD31 staining shows no continuous luminal endothelial lining (Figure 5C). Movat’s pentachrome reveals a red layer lining the lumen, consistent with fibrin deposition, which varies in thickness (measuring up to ∼220 μm). Focal mid-graft CD31-positive clusters are observed (Figure S6), indicating the onset of endothelialization at this stage. However, the relatively limited cellular infiltration and lack of substantial endothelialization of H-ENGs at two weeks contrast with our prior findings in non-heparinized ENG-based iliac artery patches that showed robust cell ingrowth and endothelialization within the same time frame. *(15)* This discrepancy likely stems from differences in graft geometry and implantation model. The tubular interposition grafts used here present a much larger luminal surface area (∼19 cm²) for cellular coverage and involve higher arterial injury, whereas patch repairs expose a broader graft-native tissue interface that facilitates more rapid migration of host cells. The limited H-ENG endothelialization at two weeks also appears inconsistent with robust in vitro proliferation, however, it likely reflects different biological contexts rather than a true contradiction. In vitro, large numbers of mature endothelial cells are seeded directly onto the graft under controlled conditions, promoting rapid proliferation and confluence. In vivo, luminal coverage depends on gradual endothelial migration from the anastomoses and incorporation of circulating progenitor cells, processes that take weeks or even months for grafts with large surface areas. Nevertheless, the observed fibrin layer likely functions as a provisional matrix for subsequent endothelial attachment and remodeling, supporting the potential for more substantial endothelialization at later time points. Despite the pilot scope, these preliminary data provide valuable insights into the early in vivo healing and remodeling processes in H-ENGs and inform the design and power calculations of more comprehensive studies.

## 3. Conclusions

This study introduces H-ENGs as a platform that integrates artery-tuned mechanics, a cell-supportive nanofibrillar architecture, and a covalently heparinized blood-material interface. While surface heparinization strategies have been reported previously, the novelty of this work lies in a one-step, PEI-enabled functionalization that preserves fiber architecture and artery-tuned nonlinear and anisotropic mechanical properties, effectively decoupling interfacial chemistry from bulk mechanics. This decoupled design supports application-tuned control of nonlinearity, anisotropy, and stiffness while retaining a stable anticoagulant interface. In vitro, H-ENGs reduce platelet adhesion and static whole-blood clot formation, display low hemolysis, and support organized endothelial monolayers without detectable cytotoxicity. In vivo pilot feasibility testing in a porcine abdominal aorta interposition model suggests that H-ENGs can be surgically implanted, withstand high-pressure arterial flow without suture hole bleeding, and retain their biomimetic mechanical properties despite tissue ingrowth and remodeling over a two-week period. XPS analysis of explanted grafts confirms the persistence of surface-bound heparin, underscoring the stability of this functionalization strategy under physiological conditions. While the current study is limited by its small sample size and short implantation period, these findings provide critical proof-of-concept evidence and a strong foundation for future preclinical studies in peripheral arterial bypass models with larger cohorts and longer follow-ups. Collectively, this work positions H-ENGs as a promising platform for next-generation synthetic vascular grafts that integrate artery-tuned mechanics, a cell-supportive microstructure, and a covalently heparinized blood-material interface.

## 4. Materials and Methods

### 4.1. Graft Manufacturing

Complete details on all chemical reagents are specified in S.1.1. ENGs were fabricated by electrospinning a polymer solution containing 8.0% (w/w) PU 82A and 2.0% (w/w) PU 55DE (10.0% total polymer concentration). To create P-ENGs, 0.5% (w/w) PEI was added to the same base formulation (10.5% total polymer concentration) to enable direct covalent coupling via 1-ethyl-3-(3-dimethylaminopropyl)carbodiimide/N-hydroxysuccinimide (EDC/NHS) bonding. All solutions were prepared in a binary solvent system of 40.0% DMF and 60.0% THF (w/w), stirred magnetically at 40 °C for at least 48 hours to ensure complete dissolution. Electrospinning was conducted using the Professional Electrospinning Device (Doxa Microfluidics) under controlled environmental conditions of 40 ± 5% relative humidity and 30 ± 2 °C. A 25-gauge, one-inch long blunt-tip needle with a 10.0 kV potential difference, a solution feed rate of 2 mL/hour, and a 10 cm needle-to-collector distance was used. Nanofibers were deposited onto an 8 mm-diameter aluminum foil-covered stainless-steel mandrel spinning at 3,000 rotations/minute. The setup included a translational stage with a 20 cm range along the collector axis, a deposition rate of 0.14 mL/cm, and a translational velocity of 6.67 cm/second. After electrospinning, all grafts were vacuum-dried overnight and released from the foil substrate. PEI incorporation was confirmed through Fourier Transform Infrared Spectroscopy (FTIR) analysis (Thermo iS50 FT-IR, Thermo-Fischer Scientific). Spectra were collected in the range of 4000–1600 cm⁻¹ to visually confirm the presence of PEI functionalization, via the peaks generated by N–H stretches of primary amine groups (Figure S1). *(100)*

Heparin (2 g/mL) was dissolved in 0.1 M 2-(N-morpholino)ethanesulfonic acid (MES) buffer (pH 5.5). To activate its carboxyl groups, EDC (3 g/mL) was added, forming reactive O-acylisourea intermediates. NHS (3 g/mL) was introduced to stabilize the intermediates. P-ENGs were immersed in 15 mL of the heparin-EDC/NHS solution at room temperature and shaken for 24 hours to enable covalent bonding, resulting in H-ENGs. H-ENGs were removed from the heparin-EDC/NHS solution and thoroughly rinsed with nanopure (18 MΩ) water to remove unbound heparin and residual reagents. Successful heparinization was characterized using a K-Alpha XPS (Thermo-Fischer Scientific) system, specifically the emergence and retention of a strong S2p peak associated with the sulfonate groups present in heparin, both after H-ENG manufacturing and again post two-week implantation.

### 4.2. Structure and Property Characterization of ENG, P-ENG, and H-ENGs

#### 4.2.1. Surgical, microstructural, and mechanical evaluation

Suture retention strength was assessed using a BioTester 5000 (CellScale) equipped with 23 N load cells (Honeywell Sensote) according to ISO 7198:2016 A.7. guidelines. Samples were tested at a crosshead speed of 100 mm/min for 45° and 90° cuts (n = 3 for all materials with three technical replicates each).

Integral water permeability (n = 3 for all materials with three technical replicates each) was evaluated according to the ISO 7198:2016 A.5.1.3. standard. Burst pressure strength testing was conducted according to ISO 7198:2016 A.5.2.2. guidelines using an Edwards Life Transducer Pressure Monitor Kit (Edwards Lifesciences). Pressures were controlled at a rate of 10 mmHg/s. Due to the highly permeable nature of the ENGs at elevated pressures, a latex tubular liner (burst pressure strength > 500 mmHg) was used to minimize leakage and enable accurate pressurization.

SEM (SNE-4500M PLUS SEC) was used to characterize ENG, P-ENG, and H-ENG microstructure and wall thickness (n = 3 for all materials with three technical replicates each) after sputter-coating the specimens (NanoImages MCM-100P) with Au for one minute. Samples were imaged under high vacuum. Graft thickness was measured (FIJI Version 2.16.0) for each graft at three different locations at 200x magnification. Images of the interior (luminal) and exterior (abluminal) surfaces of the graft were acquired at 1,000x magnification. Fiber diameter was quantified using manual measurements of a minimum of 90 fibers per image (three images per replicate) to ensure accurate comparison.

Planar biaxial tensile testing was performed using 2.5 N load cells (Honeywell Sensotec) to characterize the mechanical properties of ENGs, P-ENGs, and H-ENGs. Biaxial strain-controlled tests were performed on 11 x 11 mm samples immersed in a 37 °C 0.9% phosphate-buffered saline (PBS) bath throughout the tests. Prior to testing, an equiaxial loading protocol (1,100 mN at 3.66 mN/second) was used. Maximum circumferential and axial strains were subsequently calculated using measured rake dimensions. The maximum stretch achieved in the force-controlled test was applied in the following stretch-controlled protocol. The maximum stretch for each sample was applied at a rate of 1% stretch/second. This process was applied a total of eleven times, the initial ten cycles serving as preconditioning. The final equiaxial test was then plotted and curve-fitted using Microsoft Excel’s linest() function (Microsoft 365, 2025). Identical mechanical evaluation protocols were applied to both explanted H-ENGs and adjacent arterial tissue.

#### 4.2.2. Water Contact Angle Measurements

Water contact angle was measured using a contact angle goniometer (Ossila Ltd) for ENGs, P-ENGs, H-ENGs, and expanded polytetrafluoroethylene (ePTFE) grafts (n = 3 for all materials with three technical replicates each). Measurements were taken immediately after placing the water droplet and again after four minutes, at 62% relative humidity and 25 °C. The contact angles were analyzed using Ossila Contact Angle software version 4.2.1.

### 4.3. In Vitro Biological Compatibility Assessments

#### 4.3.1. In Vitro Hemocompatibility Assessments

For hemolysis assays, ENGs, P-ENGs, H-ENGs (n = 3 for all materials with three technical replicates each), and commercial ePTFE (n = 1 with three technical replicates) graft samples were prepared by cutting each graft into 1 cm² pieces, stabilizing in PBS for 24 hours. Fresh porcine blood collected in citrate phosphate dextrose adenine-1 (CPDA-1) anticoagulant bags (Jorgensen Laboratories) was centrifuged to isolate red blood cells (RBCs) *(101)*, which were then used to prepare a 2% RBC suspension in PBS. Each sample was incubated with 500 µL of RBC suspension in 1.5 mL microcentrifuge tubes at 37 °C for two hours. After incubation, the tubes were centrifuged at 800 × g for 12 minutes, and 200 µL of supernatant from each sample was transferred to a 96-well plate. Absorbance was measured at 545 nm using a microplate reader. Triton X-100-treated RBC suspension served as the positive control, while RBC suspension without exposure to any sample served as the negative control. The percentage of hemolysis was calculated using Equation 1, where ABS_s_ is the absorbance of the sample, ABS_p_ is the absorbance of the positive control, and ABS_n_ is the absorbance of the negative control.

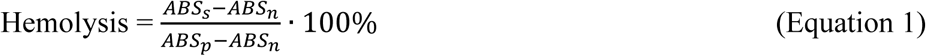

For platelet adhesion tests, ENGs, P-ENGs, H-ENGs (n = 3 per graft type with three technical replicates each), latex (n = 1 with three technical replicates), and commercial ePTFE grafts (GORE-TEX, n = 1 with three technical replicates) were used. Grafts were cut into 1 cm² samples and stabilized in PBS for 24 hours. Fresh porcine blood collected identically to the hemolysis assays was used to prepare leukocyte-rich platelet-rich plasma (PRP) *(101)*. Each sample was incubated in 500 μL leukocyte-rich PRP at 37 °C for one hour, then rinsed three times with PBS to remove non-adherent platelets. Adherent platelets were fixed by immersion in 2.5% glutaraldehyde in PBS for one hour at room temperature. The samples were then dehydrated through a graded ethanol series (30%, 50%, 70%, 90%, and 100%) for ten minutes at each concentration. Dehydrated samples were placed in a vacuum desiccator overnight to ensure complete removal of residual moisture. Samples were sputter-coated identically to 4.2.1. and imaged at 1000× magnification under high vacuum using (SNE-4500M PLUS SCE). Nine images per sample were analyzed by manual counting, and platelet adhesion was quantified in platelets/mm².

ENGs, P-ENGs, H-ENGs, commercial ePTFE grafts (GORE-TEX), and latex were cut into 1 cm² samples (n = 3 for all samples with three technical replicates each). Grafts were pre-weighed dry (W_0_) and then pre-soaked in PBS for 60 minutes at room temperature. Fresh porcine blood collected in CPDA-1 anticoagulant bags was warmed at 37 °C for ten minutes, and immediately recalcified to a final Ca²+ concentration of 10 mM by adding 0.1 M calcium chloride stock with gentle inversion. Each sample was placed in a 12-well plate with 1.0 mL of recalcified blood, covered, and incubated at 37 °C for 60 minutes. After incubation, non-adherent blood was decanted, and samples were gently rinsed three times with PBS, fixed in 2.5% glutaraldehyde in PBS for 30 minutes, and dehydrated through a graded ethanol series (identical to platelet adhesion assay). Dehydrated samples were placed in a vacuum desiccator overnight prior to re-weighing (Wı). Dry thrombus mass was calculated as Wı–Wo and reported normalized to surface area (mg/cm²).

#### 4.3.2. In Vitro Cytocompatibility Assessments

Cytotoxicity analysis of ENGs, P-ENGs, and H-ENGs (n = 3 with four technical replicates each) was conducted according to ISO 10993-5 standards. HUVECs were used for the experiments, along with an endothelial cell growth medium, which consisted of a basal medium, a supplement mix, and antibiotics (1% Pen-Strep). Two days prior to the experiments, one million cells at the fourth passage were seeded into 75 cm² tissue culture flasks and incubated at 37°C with 5% CO_2_ for approximately 48 hours or until they reached 85-95% confluence. For cell detachment, Trypsin-EDTA was diluted to a final concentration of 0.05% using Dulbecco’s PBS (full cell lines and chemical information can be found in S.1.2.). For cytotoxicity assays, 7,000 cells per well were seeded into a 96-well plate (n = 4 for each graft tested, as well as positive and negative controls) and incubated for ∼16-24 hours to allow cell attachment. Once the cells had settled, 4 mm^2^ squares of each graft treatment (ENG, P-ENG, and H-ENG) were cut and placed into wells for ∼24 hours of incubation (± 2 hours). For the positive control, 10 µL of cell lysis buffer was added one hour before the final measurements to induce maximal cell death. For the negative control, cells were cultured with the same endothelial cell complete media used previously, without exposure to test samples. Commercial ePTFE graft material was included as a reference material for comparative assessment (GORE-TEX). After incubation, the materials were removed, and the CyQUANT™ Cell Proliferation Assay, CyQUANT™ LDH Cytotoxicity Assay, and CyQUANT™ XTT Cell Viability Assay were performed according to the manufacturer’s protocols. Results were quantified using a BioTek Synergy HTX Plate Reader. According to ISO 10993-5, a reduction in cell viability by more than 30% was considered indicative of a cytotoxic effect for all three assays.

Qualitative cellular analysis was performed via immunofluorescence staining of HUVECs seeded onto ENG-type materials as well as ePTFE. Using the same cell line in the fifth passage and culture media as previously described. Graft samples were cut into 20 mm-diameter discs and placed in a 12-well plate using CellCrown inserts, with parafilm-sealed bottoms to prevent leakage. Inserts were pre-conditioned by soaking in 200 μL of complete HUVEC media at 37 °C for 24 ± 2 hours. Subsequently, 200,000 cells/cm² were seeded onto both inserts and 12-well plate surfaces (negative control), followed by 24 hours of incubation (37 °C, 5% CO2). Fixed cells (10% formalin) were permeabilized (0.2% Triton X-100), blocked (SuperBlock), and stained with p120 Antibody (G-7) Alexa Fluor® 488 and DAPI. Scaffolds were then sectioned, mounted in glycerol, and imaged using a Leica Mica microscope and LAS X Core Software (version 5.3.0).

### 4.4. In Vivo Feasibility Evaluation

#### 4.4.1. Surgical Implantation in the Porcine Abdominal Aorta Model

The acute in vivo feasibility of H-ENGs was assessed in n = 2 ∼60 kg male domestic swine (*Sus scrofa domesticus*, USDA Numbers 8506 and 8505). Animal studies were approved by the Institutional Animal Care and Use Committee (IACUC) at the University of Nebraska Medical Center (Protocol Number 23-013-04-FC). Beginning three days prior to surgery, animals were given a daily dose (81 mg) of acetylsalicylic acid (Aspirin). The animals also received 1 gram IV Cefazolin within an hour prior to beginning the procedure. Prior to implantation, H-ENGs were sterilized with ethylene oxide and soaked in heparinized phosphate buffer saline immediately prior to implantation into the abdominal aorta. Under general anesthesia, the abdominal aorta was exposed through a transabdominal incision. The animals were systemically heparinized (100-200 units/kg), and the abdominal aorta was clamped proximal and distal to the implant site. A ∼7-cm-long H-ENG was implanted onto the abdominal aorta using a continuous 5-0 polypropylene suture (Ethicon). The clamps were then removed after flushing the artery and tying down the suture line. Postoperatively, animals were given a daily dose of Aspirin for two weeks. Graft patency and thrombus accumulation were assessed with duplex ultrasound examinations immediately after implantation and two weeks post-implantation. After two weeks, animals were euthanized, and the abdominal aorta and graft was collected for visual inspection, mechanical testing, and histological assessment.

#### 4.4.2. Duplex Ultrasound Evaluations

Duplex ultrasound of the native abdominal aorta and the implanted H-ENG were taken both immediately after implantation and two weeks post-implantation using the Philips CX50 Portable Ultrasound System equipped with Philips L12-3 and L15-7io transducers. The level of graft compliance was visually identifiable during the surgery. Cross-sectional view duplex ultrasound videos from both implantation and two-week ultrasounds were tracked (Kinovea 2023.1.2) to identify diastolic and systolic time points for both the H-ENG and the abdominal aorta (Figure 4B). The perimeter at each of these time points was then determined, and pulsatility was calculated for five pulses for both the native abdominal aorta and the H-ENG (**Equation 2**) *(102)*.

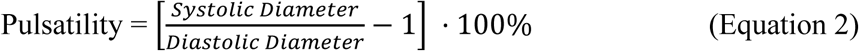

### 4.5. Ex Vivo Feasibility Evaluation

#### 4.5.1. Evaluation of Mechanical Properties Post-Implantation

Mechanical testing post-implantation was performed identically to section 4.2.1. Samples were tested within 24 hours post-explantation to preserve the mechanical properties of any biological tissues in or surrounding the H-ENG.

#### 4.5.2. Verification of Surface Heparin Presence After Two Weeks In Vivo

XPS Evaluation post-implantation was performed identically to section 4.1.1.

#### 4.5.3. Early Remodeling Evaluation Through Histological Imaging

Following necropsy, H-ENGs and adjacent aortic segments were excised and collected. Cross-sectional rings of the native abdominal aorta and H-ENG graft mid-section and longitudinal sections of abdominal aorta/H-ENG distal anastomoses were cut and fixed in 10% phosphate-buffered formalin for 24-48 hours. Following fixation, all samples were transferred to 70% ethanol solution prior to paraffin embedding. The embedded samples were cut into 5 µm-thick sections and stained with H&E, α-SMA, CD31, and Movat’s Pentachrome stains. The stained slides were then scanned at 20x magnification and converted to .jpg image file format.

### 4.6. Statistical Analysis

#### 4.6.1. Bayesian Statistics

All descriptive statistics were performed in Microsoft Excel (Microsoft 365, 2025) using the average() and stdev.s() functions. A Bayesian multilevel regression model was implemented in R Studio (Mariposa Orchid, May 2025) to investigate the effect of treatment on fabric properties, accounting for variations across individual fabrics. Fabric identification number was included as a random effect, while treatment was specified as a fixed effect. The dependent variables: circumferential stiffness, axial stiffness, and graft thickness were analyzed using a multivariate model. Water contact angle, fiber diameter, and platelet adhesion were analyzed using univariate models.

For each model, four Markov Chain Monte Carlo chains were run, with 2000 iterations per chain. To ensure reliable convergence and reduce divergent transitions, the target average acceptance probability was set to 0.95. All models were assessed for convergence using standard diagnostic plots and statistics (R-hat values, effective sample size (ESS), bulk and tail EES, and posterior predictive checks).

Given that some datasets exhibited non-normal distribution, two primary approaches were employed using the mvbind() and brms() functions. Initially, a Gaussian distribution was defined, allowing for a comparison with traditional linear mixed models. Subsequently, to better accommodate non-normal data, an alternative Poisson distribution was also fitted. These two fits were then compared using the loo() function and the better fit was used.

#### 4.6.2. T-tests

All descriptive statistics were performed in Microsoft Excel (Microsoft 365, 2025) using the average() and stdev.s() functions. For tests with low sample size (n = 2), normality and homogeneity of variance were not checked, as tests are not valid for such small sample sizes. Homoscedastic t-tests were performed using Microsoft Excel (Microsoft 365, 2025) t.test() function, where α = .05.

## Supporting Information

Complete reagents and materials are listed; FTIR spectra confirm PEI incorporation through N–H/O–H stretching peaks; mechanical testing shows consistent suture retention and sufficient burst pressure capacity; H-ENG water contact angle decreases from ∼90° to ∼0° within 4 minutes demonstrating complete wetting; SEM reveals high platelet/WBC adhesion on P-ENG but minimal adhesion on H-ENG and ePTFE; HUVEC imaging shows robust cell attachment across graft types; histology from porcine implants shows preserved graft structure with localized thrombus only near one anastomosis; ultrasound confirms maintained pulsatility and flow at proximal and distal anastomoses; tables report microstructural/mechanical properties, hemolysis/platelet adhesion/clotting metrics, cytotoxicity assay results, and duplex imaging assessments.

## Data Availability Statement

Data will be made available upon request. Full histology images are openly available in Figshare at https://doi.org/10.6084/m9.figshare.30179797, reference number 30179797.

## Author Contributions

**Elizabeth Zermeno:** Data curation, Formal analysis, Investigation, Methodology, Visualization, Writing – original draft, Writing – review & editing. **John Kapitan:** Data curation, Investigation, Methodology, Writing – original draft, Writing – review & editing. **Alexander Sandquist:** Data curation, Investigation, Methodology, Writing – original draft, Writing – review & editing. **Amanda Reke:** Investigation, Methodology, Visualization, Writing – original draft, Writing – review & editing. **Apurbo Kumar Paul:** Data curation, Investigation, Methodology, Visualization, Writing – original draft. **Jason MacTaggart:** Conceptualization, Investigation, Methodology, Project administration, Resources, Supervision, Writing – review & editing. **Stephen Morin:** Conceptualization, Funding acquisition, Investigation, Project administration, Writing – original draft, Writing – review & editing. **Kaspars Maleckis:** Conceptualization, Funding acquisition, Investigation, Project administration, Resources, Supervision, Visualization, Writing – original draft, Writing – review & editing.

## Declaration of Generative AI

During the preparation of this work the author(s) used ChatGPT 5.0 and Gemini 3 Pro in order to screen for grammatical errors. After using this tool/service, the author(s) reviewed and edited the content as needed and take(s) full responsibility for the content of the published article.

## References

1. Martin, S. S.; Aday, A. W.; Almarzooq, Z. I.; Anderson, C. A. M.; Arora, P.; Avery, C. L.;, et al. 2024 Heart Disease and Stroke Statistics: A Report of US and Global Data From the American Heart Association. Circulation 2024, 149(8). doi:10.1161/CIR.0000000000001209.

2. Szafron, J. M.; Heng, E. E.; Boyd, J.; Humphrey, J. D.; Marsden, A. L. Hemodynamics and Wall Mechanics of Vascular Graft Failure. *Arteriosclerosis*, Thrombosis, and Vascular Biology 2024, 44(5), 1065–1085. doi:10.1161/ATVBAHA.123.318239.

3. Abbott, W. M.; Megerman, J.; Hasson, J. E.; L’Italien, G.; Warnock, D. F. Effect of compliance mismatch on vascular graft patency. Journal of Vascular Surgery 1987, 5(2), 376–382. doi:10.1016/0741-5214(87)90148-0.

4. Jacot, J. G.; Wong, J. Y. Endothelial injury induces vascular smooth muscle cell proliferation in highly localized regions of a direct contact co-culture system. Cell biochemistry and biophysics 2008, 52(1), 37–46. doi:10.1007/s12013-008-9023-6.

5. Giudiceandrea, A.; Seifalian, A. M.; Krijgsman, B.; Hamilton, G. Effect of prolonged pulsatile shear stress in vitro on endothelial cell seeded PTFE and compliant polyurethane vascular grafts. Eur J Vasc Endovasc Surg 1998, 15(2), 147–154. doi:10.1016/S1078-5884(98)80136-6.

6. Kamenskiy, A. V.; Dzenis, Y. A.; Kazmi, S. A. J.; Pemberton, M. A.; Pipinos, I. I. I.; Phillips, N. Y.;, et al. Biaxial mechanical properties of the human thoracic and abdominal aorta, common carotid, subclavian, renal and common iliac arteries. Biomechanics and modeling in mechanobiology 2014, 13(6), 1341–1359. doi:10.1007/s10237-014-0576-6.

7. Hu, K.; Li, Y.; Ke, Z.; Yang, H.; Lu, C.; Li, Y.;, et al. History, progress and future challenges of artificial blood vessels: a narrative review. Biomaterials Translational 2022, 3(1), 81–98. doi:10.12336/biomatertransl.2022.01.008.

8. Vascular Grafts Market Size, Share & Growth Report, 2030; 2025. <https://www.grandviewresearch.com/industry-analysis/vascular-graft-market#> Accessed 25.03.15.

9. Tai, N. R.; Salacinski, H. J.; Edwards, A.; Hamilton, G.; Seifalian, A. M. Compliance properties of conduits used in vascular reconstruction. British Journal of Surgery 2000, 87(11), 1516–1524. doi:10.1046/j.1365-2168.2000.01566.x.

10. Nezarati, R. M.; Eifert, M. B.; Dempsey, D. K.; Cosgriff-Hernandez, E. Electrospun vascular grafts with improved compliance matching to native vessels. Journal of Biomedical Materials Research Part B: Applied Biomaterials 2015, 103(2), 313–323. doi:10.1002/jbm.b.33201.

11. Qiu, S.; Du, J.; Zhu, T.; Zhang, H.; Chen, S.; Wang, C.;, et al. Electrospun compliant heparinized elastic vascular graft for improving the patency after implantation. International Journal of Biological Macromolecules 2023, 253, 126598. doi:10.1016/j.ijbiomac.2023.126598.

12. Anttila, E.; Balzani, D.; Desyatova, A.; Deegan, P.; MacTaggart, J.; Kamenskiy, A. Mechanical damage characterization in human femoropopliteal arteries of different ages. Acta Biomaterialia 2019, 90, 225–240. doi:10.1016/j.actbio.2019.03.053.

13. Jadidi, M.; Habibnezhad, M.; Anttila, E.; Maleckis, K.; Desyatova, A.; MacTaggart, J.;, et al. Mechanical and structural changes in human thoracic aortas with age. Acta Biomaterialia 2020, 103, 172–188. doi:10.1016/j.actbio.2019.12.024.

14. Trubel, W.; Schima, H.; Moritz, A.; Raderer, F.; Windisch, A.; Ullrich, R.;, et al. Compliance mismatch and formation of distal anastomotic intimal hyperplasia in externally stiffened and lumen-adapted venous grafts. Eur J Vasc Endovasc 1995, 10(4), 415–423.

15. Maleckis, K.; Kamenskiy, A.; Lichter, E. Z.; Oberley-Deegan, R.; Dzenis, Y.; MacTaggart, J. Mechanically tuned vascular graft demonstrates rapid endothelialization and integration into the porcine iliac artery wall. Acta Biomaterialia 2021, 125, 126–137. doi:10.1016/j.actbio.2021.01.047.

16. Humphrey, J. D.; Schwartz, M. A.; Tellides, G.; Milewicz, D. M. Role of Mechanotransduction in Vascular Biology. Circulation Research 2015, 116(8), 1448–1461. doi:10.1161/CIRCRESAHA.114.304936.

17. Zhang, Y.; Li, X. S.; Guex, A. G.; Liu, S. S.; Müller, E.; Malini, R. I.;, et al. A compliant and biomimetic three-layered vascular graft for small blood vessels. Biofabrication 2017, 9(2), 025010. doi:10.1088/1758-5090/aa6bae.

18. Spencer, M.; Denison, A. In Handbook of Physiology; Am. Physiol. Soc: Washington, DC, 1963; Vol. II, p 842.

19. Chiu, J.-J.; Chien, S. Effects of disturbed flow on vascular endothelium: pathophysiological basis and clinical perspectives. Physiological Reviews 2011, 91(1), 327–387. doi:10.1152/physrev.00047.2009.

20. Fathi-Karkan, S.; Banimohamad-Shotorbani, B.; Saghati, S.; Rahbarghazi, R.; Davaran, S. A critical review of fibrous polyurethane-based vascular tissue engineering scaffolds. Journal of Biological Engineering 2022, 16(1), 6. doi:10.1186/s13036-022-00286-9.

21. Okoshi, T.; Soldani, G.; Goddard, M.; Galletti, P. M. Very small-diameter polyurethane vascular prostheses with rapid endothelialization for coronary artery bypass grafting. The Journal of Thoracic and Cardiovascular Surgery 1993, 105(5), 791–795.

22. Ozdemir, S.; Yalcin-Enis, I.; Yalcinkaya, B.; Yalcinkaya, F. An Investigation of the Constructional Design Components Affecting the Mechanical Response and Cellular Activity of Electrospun Vascular Grafts. Membranes 2022, 12(10), 929. doi:10.3390/membranes12100929.

23. Narayan, D.; Venkatraman, S. S. Effect of pore size and interpore distance on endothelial cell growth on polymers. Journal of Biomedical Materials Research. Part A 2008, 87(3), 710–718. doi:10.1002/jbm.a.31749.

24. Mukasheva, F.; Adilova, L.; Dyussenbinov, A.; Yernaimanova, B.; Abilev, M.; Akilbekova, D. Optimizing scaffold pore size for tissue engineering: insights across various tissue types. Frontiers in Bioengineering and Biotechnology 2024, 12, 1444986. doi:10.3389/fbioe.2024.1444986.

25. Clowes, A. W. Mechanisms of Arterial Graft Healing. 1986, 123(2).

26. Golden, M. A.; Hanson, S. R.; Kirkman, T. R.; Schneider, P. A.; Clowes, A. W. Healing of polytetrafluoroethylene arterial grafts is influenced by graft porosity. Journal of Vascular Surgery 1990, 11(6), 838–845. doi:10.1016/0741-5214(90)90082-L.

27. Zhang, Z.; Wang, Z.; Liu, S.; Kodama, M. Pore size, tissue ingrowth, and endothelialization of small-diameter microporous polyurethane vascular prostheses. Biomaterials 2004, 25(1), 177–187. doi:10.1016/S0142-9612(03)00478-2.

28. Bosiers, M.; Deloose, K.; Verbist, J.; Schroë, H.; Lauwers, G.; Lansink, W.;, et al. Heparin-bonded expanded polytetrafluoroethylene vascular graft for femoropopliteal and femorocrural bypass grafting: 1-year results. Journal of Vascular Surgery 2006, 43(2), 313–318. doi:10.1016/j.jvs.2005.10.037.

29. Ratner, B. Vascular Grafts: Technology Success/Technology Failure. BME Frontiers 4, 0003. doi:10.34133/bmef.0003.

30. Holubec, H.; Hunter, G.; Chvapil, M.; Chvapil, T.; Bernhard, V.; Misiorowski, R.;, et al. The Relationship Between PTFE Graft Ultrastructure and Cellular Ingrowth: The Influence of an Autologous Jugular Vein Wrap. In Biomaterials’ Mechanical Properties. Kambic, H., Yokobori, A., Jr, Eds.; ASTM International, 1994; Vol. STP1173-EB, p 0. doi:10.1520/STP18093S.

31. Polyethylene Terephthalate - an overview | ScienceDirect Topics. <https://www.sciencedirect.com/topics/pharmacology-toxicology-and-pharmaceutical-science/polyethylene-terephthalate> Accessed 25.07.17.

32. Kondo, Y.; Muto, A.; Dardik, A.; Nishibe, M.; Nishibe, T. Perigraft Seroma After Surgical Aortoiliac Aneurysm Repair with Knitted Polyester Grafts: Report of Two Cases. Annals of Vascular Diseases 2009, 2(1), 44–46. doi:10.3400/avd.AVDcr08010.

33. Lu, P. G.; Erben, Y.; Sheaffer, W. W.; Pierce, A. T.; Mendes, B.; DeMartino, R.;, et al. “Large Diameter” Aortic Endografts are Associated With Aneurysm Sac Expansion. Annals of Vascular Surgery 2022, 87, 225–230. doi:10.1016/j.avsg.2022.04.046.

34. Matsushita, A.; Tsunoda, Y.; Hattori, T.; Mihara, W. Late Disruption of a Polyethylene Terephthalate Aortic Graft 30 Years after Initial Graft Placement. EJVES Short Reports 2017, 38, 4–7. doi:10.1016/j.ejvssr.2017.11.001.

35. Yener, A.; Yener, N. Serous fluid leakage without seroma after aortobifemora bypass operation. Annals of Thoracic and Cardiovascular Surgery: Official Journal of the Association of Thoracic and Cardiovascular Surgeons of Asia 2002, 8(1), 54–55.

36. Matsuzaki, Y.; Iwaki, R.; Reinhardt, J. W.; Chang, Y.-C.; Miyamoto, S.; Kelly, J.;, et al. The effect of pore diameter on neo-tissue formation in electrospun biodegradable tissue-engineered arterial grafts in a large animal model. Acta Biomaterialia 2020, 115, 176–184. doi:10.1016/j.actbio.2020.08.011.

37. Wu, J.; Hong, Y. Enhancing cell infiltration of electrospun fibrous scaffolds in tissue regeneration. Bioactive Materials 2016, 1(1), 56–64. doi:10.1016/j.bioactmat.2016.07.001.

38. Ameer, J. M.; Pr, A. K.; Kasoju, N. Strategies to Tune Electrospun Scaffold Porosity for Effective Cell Response in Tissue Engineering. Journal of Functional Biomaterials 2019, 10(3), 30. doi:10.3390/jfb10030030.

39. Zhang, X.; Sun, Q.; Liang, X.; Gu, P.; Hu, Z.; Yang, X.;, et al. Stretchable and negative-Poisson-ratio porous metamaterials. Nature Communications 2024, 15, 392. doi:10.1038/s41467-024-44707-3.

40. Sawadkar, P.; Mandakhbayar, N.; Patel, K. D.; Buitrago, J. O.; Kim, T. H.; Rajasekar, P.;, et al. Three dimensional porous scaffolds derived from collagen, elastin and fibrin proteins orchestrate adipose tissue regeneration. Journal of Tissue Engineering 2021, 12, 20417314211019238. doi:10.1177/20417314211019238.

41. Fischer, T.; Hayn, A.; Mierke, C. T. Fast and reliable advanced two-step pore-size analysis of biomimetic 3D extracellular matrix scaffolds. Scientific Reports 2019, 9(1), 8352. doi:10.1038/s41598-019-44764-5.

42. Harik, L.; Perezgrovas-Olaria, R.; Soletti, G.; Dimagli, A.; Alzghari, T.; An, K. R.;, et al. Graft thrombosis after coronary artery bypass surgery and current practice for prevention. Frontiers in Cardiovascular Medicine 2023, 10, 1125126. doi:10.3389/fcvm.2023.1125126.

43. Gupta, S.; Puttaiahgowda, Y. M.; Deiglmayr, L. Recent advances in the design and immobilization of heparin for biomedical application: A review. International Journal of Biological Macromolecules 2024, 264, 130743. doi:10.1016/j.ijbiomac.2024.130743.

44. Zhu, T.; Gu, H.; Zhang, H.; Wang, H.; Xia, H.; Mo, X.;, et al. Covalent grafting of PEG and heparin improves biological performance of electrospun vascular grafts for carotid artery replacement. Acta Biomaterialia 2021, 119, 211–224. doi:10.1016/j.actbio.2020.11.013.

45. Li, C.; Yang, Q.; Chen, D.; Zhu, H.; Chen, J.; Liu, R.; et al. Polyethyleneimine-assisted co-deposition of polydopamine coating with enhanced stability and efficient secondary modification. 2022. doi:10.1039/D2RA05130C.

46. Kim, C. H.; Kim, Y.; Karna, S.; Yoo, S. M.; Lee, J. H.; Kim, Y. J.;, et al. Three-dimensional customized artificial grafts functionalized with biomimetic softness and anticoagulant heparin-dopamine surface modification: Preclinical study for practical applications. International Journal of Biological Macromolecules 2025, 299, 140002. doi:10.1016/j.ijbiomac.2025.140002.

47. Tabish, T. A.; Thorat, N. D.; Narayan, R. J. Mechanical behaviour of nitric oxide releasing polymers for cardiovascular bypass grafts. Mechanics of Materials 2023, 176, 104520. doi:10.1016/j.mechmat.2022.104520.

48. Ott, D.; Pulfer, S.; Smith, D. Incorporation of Nitric Oxide-Releasing Crosslinked Polyethyleneimine Microspheres into Vascular Grafts. Journal of Biomedical Materials Research 1997, 37(2), 182–189.

49. Allen, B. T.; Long, J. A.; Clark, R. E.; Sicard, G. A.; Hopkins, K. T.; Welch, M. J. Influence of endothelial cell seeding on platelet deposition and patency in small-diameter Dacron arterial grafts. Journal of Vascular Surgery 1984, 1(1), 224–233. doi:10.1016/0741-5214(84)90201-5.

50. Noishiki, Y.; Yamane, Y.; Tomizawa, Y.; Okoshi, T.; Satoh, S.; Wildevuur, C. R. H.;, et al. Rapid endothelialization of vascular prostheses by seeding autologous venous tissue fragments. The Journal of Thoracic and Cardiovascular Surgery 1992, 104(3), 770–778. doi:10.1016/S0022-5223(19)34749-X.

51. Peng, G.; Yao, D.; Niu, Y.; Liu, H.; Fan, Y. Surface Modification of Multiple Bioactive Peptides to Improve Endothelialization of Vascular Grafts. Macromolecular Bioscience 2019, 19(5), e1800368. doi:10.1002/mabi.201800368.

52. Lazarides, M. K.; Argyriou, C.; Antoniou, G. A.; Georgakarakos, E.; Georgiadis, G. S. Lack of evidence for use of heparin-bonded grafts in access surgery: a meta-analysis. Seminars in Vascular Surgery 2016, 29(4), 192–197. doi:10.1053/j.semvascsurg.2016.08.003.

53. Lindholt, J. S.; Houlind, K.; Gottschalksen, B.; Pedersen, C. N.; Ravn, H.; Viddal, B.;, et al. Five-year outcomes following a randomized trial of femorofemoral and femoropopliteal bypass grafting with heparin-bonded or standard polytetrafluoroethylene grafts. The British Journal of Surgery 2016, 103(10), 1300–1305. doi:10.1002/bjs.10246.

54. Fischer, M. J. E. Amine coupling through EDC/NHS: a practical approach. *Methods in Molecular Biology (Clifton*, N.J.) 2010, 627, 55–73. doi:10.1007/978-1-60761-670-2_3.

55. Shanthi, P.; Hanumantha, P.; Ramalinga, K.; Gattu, B.; Datta, M.; Kumta, P. Sulfonic Acid Based Complex Framework Materials (CFM): Nanostructured Polysulfide Immobilization Systems for Rechargeable Lithium–Sulfur Battery. Journal of The Electrochemical Society 2019, 166, A1827–A1835. doi:10.1149/2.0251910jes.

56. Matsuzaki, Y.; Ulziibayar, A.; Shoji, T.; Shinoka, T. Heparin-Eluting Tissue-Engineered Bioabsorbable Vascular Grafts. Applied Sciences 2021, 11(10), 4563. doi:10.3390/app11104563.

57. Aslani, S.; Kabiri, M.; HosseinZadeh, S.; Hanaee-Ahvaz, H.; Taherzadeh, E. S.; Soleimani, M. The applications of heparin in vascular tissue engineering. Microvascular Research 2020, 131, 104027. doi:10.1016/j.mvr.2020.104027.

58. Heyligers, J. M. M.; Verhagen, H. J. M.; Rotmans, J. I.; Weeterings, C.; de Groot, P. G.; Moll, F. L.;, et al. Heparin immobilization reduces thrombogenicity of small-caliber expanded polytetrafluoroethylene grafts. Journal of Vascular Surgery 2006, 43(3), 587–591. doi:10.1016/j.jvs.2005.10.038.

59. Nie, C.; Ma, L.; Cheng, C.; Deng, J.; Zhao, C. Nanofibrous heparin and heparin-mimicking multilayers as highly effective endothelialization and antithrombogenic coatings. Biomacromolecules 2015, 16(3), 992–1001. doi:10.1021/bm501882b.

60. Qiu, X.; Lee, B. L.-P.; Ning, X.; Murthy, N.; Dong, N.; Li, S. End-point immobilization of heparin on plasma-treated surface of electrospun polycarbonate-urethane vascular graft. Acta Biomaterialia 2017, 51, 138–147. doi:10.1016/j.actbio.2017.01.012.

61. Cheng, C.; Sun, S.; Zhao, C. Progress in heparin and heparin-like/mimicking polymer-functionalized biomedical membranes. 2014. doi:10.1039/C4TB01390E.

62. The suture retention test, revisited and revised. ResearchGate. <https://www.researchgate.net/publication/319283670_The_suture_retention_test_revisited_and_revised> Accessed 25.10.10.

63. Madhavan, K.; Elliott, W. H.; Bonani, W.; Monnet, E.; Tan, W. Mechanical and biocompatible characterizations of a readily available multilayer vascular graft. Journal of Biomedical Materials Research Part B: Applied Biomaterials 2013, 101B(4), 506–519. doi:10.1002/jbm.b.32851.

64. Greenhalgh, E. S.; Dunn, M. W. Modeling Blood Flow Through Vascular Grafts.

65. Palatini, P.; Mos, L.; Munari, L.; Valle, F.; Del Torre, M.; Rossi, A.;, et al. Blood pressure changes during heavy-resistance exercise. Journal of Hypertension. Supplement: Official Journal of the International Society of Hypertension 1989, 7(6), S72–73. doi:10.1097/00004872-198900076-00032.

66. Suroto, N. S.; Al Fauzi, A.; Widiyanti, P.; Bella, F. R. Biocompatibility evaluation of electrospun Poly-L lactic Acid-chitosan immobilized with heparin as scaffold for vascular tissue repair. Journal of Science: Advanced Materials and Devices 2023, 8(3), 100594. doi:10.1016/j.jsamd.2023.100594.

67. Liu, P.; Liu, X.; Yang, L.; Qian, Y.; Lu, Q.; Shi, A.;, et al. Enhanced hemocompatibility and rapid magnetic anastomosis of electrospun small-diameter artificial vascular grafts. Frontiers in Bioengineering and Biotechnology 2024, 12. doi:10.3389/fbioe.2024.1331078.

68. Vogler, E. A. Structure and reactivity of water at biomaterial surfaces. Advances in Colloid and Interface Science 1998, 74(1), 69–117. doi:10.1016/S0001-8686(97)00040-7.

69. Correlation of proliferation, morphology and biological responses of fibroblasts on LDPE with different surface wettability: Journal of Biomaterials Science, Polymer Edition: Vol 18, No 5. <https://www.tandfonline.com/doi/abs/10.1163/156856207780852514> Accessed 25.07.09.

70. Lee, J. H.; Lee, S. J.; Khang, G.; Lee, H. B. The Effect of Fluid Shear Stress on Endothelial Cell Adhesiveness to Polymer Surfaces with Wettability Gradient. Journal of Colloid and Interface Science 2000, 230(1), 84–90. doi:10.1006/jcis.2000.7080.

71. Davoudi, P.; Assadpour, S.; Derakhshan, M. A.; Ai, J.; Solouk, A.; Ghanbari, H. Biomimetic modification of polyurethane-based nanofibrous vascular grafts: A promising approach towards stable endothelial lining. Materials Science and Engineering: C 2017, 80, 213–221. doi:10.1016/j.msec.2017.05.140.

72. Niemczyk-Soczynska, B.; Gradys, A.; Sajkiewicz, P. Hydrophilic Surface Functionalization of Electrospun Nanofibrous Scaffolds in Tissue Engineering. Polymers 2020, 12(11), 2636. doi:10.3390/polym12112636.

73. Miyake, J. Cellulose Proton Conductor: Both Sulfonic Acid and Hydrophobic Group Functionalization Enable High Proton Conductivity. JACS Au 2025. doi:10.1021/jacsau.5c00547.

74. Zhang, J.; Guo, J.; Zhang, J.; Li, D.; Zhong, M.; Gu, Y.;, et al. Heparin and Gelatin Co-Functionalized Polyurethane Artificial Blood Vessel for Improving Anticoagulation and Biocompatibility. Bioengineering 2025, 12(3), 304. doi:10.3390/bioengineering12030304.

75. Xu, L.-C.; Bauer, J. W.; Siedlecki, C. A. Proteins, platelets, and blood coagulation at biomaterial interfaces. Colloids and Surfaces B: Biointerfaces 2014, 124, 49–68. doi:10.1016/j.colsurfb.2014.09.040.

76. Tzoneva, R.; Faucheux, N.; Groth, T. Wettability of substrata controls cell–substrate and cell–cell adhesions. Biochimica et Biophysica Acta (BBA) - General Subjects 2007, 1770(11), 1538–1547. doi:10.1016/j.bbagen.2007.07.008.

77. Altshuler, P.; Nahirniak, P.; Welle, N. J. Saphenous Vein Grafts. In StatPearls; StatPearls Publishing: Treasure Island (FL), 2025.

78. Naoum, J. J.; Arbid, E. J. Bypass Surgery in Limb Salvage: Polytetrafluoroethylene Prosthetic Bypass. Methodist DeBakey Cardiovascular Journal 2012, 8(4), 43–46. doi:10.14797/mdcj-8-4-43.

79. Tulloch, A. W.; Chun, Y.; Levi, D. S.; Mohanchandra, K. P.; Carman, G. P.; Lawrence, P. F.;, et al. Super Hydrophilic Thin Film Nitinol Demonstrates Reduced Platelet Adhesion Compared with Commercially Available Endograft Materials. Journal of Surgical Research 2011, 171(1), 317–322. doi:10.1016/j.jss.2010.01.014.

80. Wang, S.; Gupta, A. S.; Sagnella, S.; Barendt, P. M.; Kottke-Marchant, K.; Marchant, R. E. Biomimetic Fluorocarbon Surfactant Polymers Reduce Platelet Adhesion on PTFE/ePTFE Surfaces. Journal of biomaterials science. Polymer edition 2009, 20(5–6), 619–635. doi:10.1163/156856209X426439.

81. Chuang, T.-W.; Masters, K. S. Regulation of polyurethane hemocompatibility and endothelialization by tethered hyaluronic acid oligosaccharides. Biomaterials 2009, 30(29), 5341–5351. doi:10.1016/j.biomaterials.2009.06.029.

82. Tan, Y.; Shao, Z.-B.; Yu, L.-X.; Xu, Y.-J.; Rao, W.-H.; Chen, L.;, et al. Polyethyleneimine modified ammonium polyphosphate toward polyamine-hardener for epoxy resin: Thermal stability, flame retardance and smoke suppression. Polymer Degradation and Stability 2016, 131, 62–70. doi:10.1016/j.polymdegradstab.2016.07.004.

83. Hautmann, A.; Kedilaya, D.; Stojanovic, S.; Radenković, M.; Marx, C.; Najman, S.;, et al. Free-standing multilayer films as growth factor reservoirs for future wound dressing applications. Biomaterials Advances 2022, 142, 213166. doi:10.1016/j.bioadv.2022.213166.

84. Miyamoto, M.; Sasakawa, S.; Ozawa, T.; Kawaguchi, H.; Ohtsuka, Y. Mechanisms of blood coagulation induced by latex particles and the roles of blood cells. Biomaterials 1990, 11(6), 385–388. doi:10.1016/0142-9612(90)90091-4.

85. Johno, H.; Takahashi, S.; Kitamura, M. Influences of Acidic Conditions on Formazan Assay: A Cautionary Note. Applied Biochemistry and Biotechnology 2010, 162(6), 1529– 1535. doi:10.1007/s12010-010-8934-z.

86. Schulz, C.; von Rüsten-Lange, M.; Krüger, A.; Lendlein, A.; Jung, F. Viability and function of primary human endothelial cells on smooth poly(ether imide) films. Clinical Hemorheology and Microcirculation 2012, 52(2–4), 267–282. doi:10.3233/CH-2012-1604.

87. Wang, D.; Xu, Y.; Wang, L.; Wang, X.; Ren, C.; Zhang, B.;, et al. Expanded Poly(tetrafluoroethylene) Blood Vessel Grafts with Embedded Reactive Oxygen Species (ROS)-Responsive Antithrombogenic Drug for Elimination of Thrombosis. ACS Applied Materials & Interfaces 2020, 12(26), 29844–29853. doi:10.1021/acsami.0c07868.

88. Zhang, Q.; Wang, C.; Babukutty, Y.; Ohyama, T.; Kogoma, M.; Kodama, M. Biocompatibility evaluation of ePTFE membrane modified with PEG in atmospheric pressure glow discharge. Journal of Biomedical Materials Research 2002, 60(3), 502–509. doi:10.1002/jbm.1294.

89. Krüger-Genge, A.; Dietze, S.; Yan, W.; Liu, Y.; Fang, L.; Kratz, K.;, et al. Endothelial cell migration, adhesion and proliferation on different polymeric substrates. Clinical Hemorheology and Microcirculation 2019, 70(4), 511–529. doi:10.3233/CH-189317.

90. Piña, R.; Santos-Díaz, A. I.; Orta-Salazar, E.; Aguilar-Vazquez, A. R.; Mantellero, C. A.; Acosta-Galeana, I.;, et al. Ten Approaches That Improve Immunostaining: A Review of the Latest Advances for the Optimization of Immunofluorescence. International Journal of Molecular Sciences 2022, 23(3), 1426. doi:10.3390/ijms23031426.

91. Simón-Polo, E.; Catalá-Ripoll, J. V.; Monsalve-Naharro, J. Á.; Gerónimo-Pardo, M. Cardiac output and the pharmacology of general anesthetics: a narrative review. Colombian Journal of Anesthesiology 2023, 51(4). doi:10.5554/22562087.e1074.

92. Lumsden, A. B.; Morrissey, N. J.; Comparison of Safety and Primary Patency Between the FUSION BIOLINE Heparin-Coated Vascular Graft and EXXCEL Soft ePTFE (FINEST) Trial Co-investigators. Randomized controlled trial comparing the safety and efficacy between the FUSION BIOLINE heparin-coated vascular graft and the standard expanded polytetrafluoroethylene graft for femoropopliteal bypass. Journal of Vascular Surgery 2015, 61(3), 703–712.e1. doi:10.1016/j.jvs.2014.10.008.

93. Miller, C. M.; Sangiolo, P.; Jacobson, J. H. Reduced anastomotic bleeding using new sutures with a needle-suture diameter ratio of one. Surgery 1987, 101(2), 156–160.

94. Heparin coatings for improving blood compatibility of medical devices. ResearchGate. <https://www.researchgate.net/publication/311977281_Heparin_coatings_for_improving_blood_compatibility_of_medical_devices> Accessed 25.10.10.

95. Hinds, M. T.; Rowe, R. C.; Ren, Z.; Teach, J.; Wu, P.; Kirkpatrick, S. J.;, et al. Development of a reinforced porcine elastin composite vascular scaffold. Journal of Biomedical Materials Research Part A 2006, 77A(3), 458–469. doi:10.1002/jbm.a.30571.

96. Hao, D.; Lin, J.; Liu, R.; Pivetti, C.; Yamashiro, K.; Schutzman, L. M.;, et al. A bio-instructive parylene-based conformal coating suppresses thrombosis and intimal hyperplasia of implantable vascular devices. Bioactive Materials 2023, 28, 467–479. doi:10.1016/j.bioactmat.2023.06.014.

97. Szafron, J. M.; Heng, E. E.; Boyd, J.; Humphrey, J. D.; Marsden, A. L. Hemodynamics and Wall Mechanics of Vascular Graft Failure. *Arteriosclerosis*, Thrombosis, and Vascular Biology 2024, 44(5), 1065–1085. doi:10.1161/ATVBAHA.123.318239.

98. Kurmaev, E. Z.; Fedorenko, V. V.; Galakhov, V. R.; Bartkowski, S.; Uhlenbrock, S.; Neumann, M.;, et al. Analysis of oxyanion (BO33-, CO32-, SO42-, PO43-, SeO44-) substitution in Y123 compounds studied by X-ray photoelectron spectroscopy. Journal of Superconductivity 1996, 9(1), 97–100. doi:10.1007/BF00728433.

99. Ha, F.; Wu, Y.; Wang, H.; Wang, T. The Reference Intervals of Whole Blood Copper, Zinc, Calcium, Magnesium, and Iron in Infants Under 1 Year Old. Biological Trace Element Research 2022, 200(1), 1–12. doi:10.1007/s12011-021-02620-6.

100. John McMurry, P. E. 24.10 Spectroscopy of Amines - Organic Chemistry | OpenStax. <https://openstax.org/books/organic-chemistry/pages/24-10-spectroscopy-of-amines> Accessed 25.06.16.

101. A Simple Double-Spin Closed Method for Preparing Platelet-Rich Plasma | Cureus. <https://www.cureus.com/articles/81542-a-simple-double-spin-closed-method-for-preparing-platelet-rich-plasma#!/> Accessed 25.04.11.

102. Zhu, W.; Wang, Y.; Chen, Y.; Liu, J.; Zhou, C.; Shi, Q.;, et al. Dynamic Changes in the Aorta During the Cardiac Cycle Analyzed by ECG-Gated Computed Tomography. Frontiers in Cardiovascular Medicine 2022, 9, 793722. doi:10.3389/fcvm.2022.793722.

