## Supporting Information for "Heparinized Elastomeric Nanofibrillar Grafts: A Novel Approach for Mechanically Tunable, Cell-Supportive, and Thromboresistant Vascular Substitutes"

### S.1. Reagents and Materials

#### S.1.1. Graft Manufacturing

Biomedical grade polyether urethanes Pellethane® 5863–82A (PU 82A) and Pellethane® 2363–55DE (PU 55DE) were purchased from Lubrizol Life Science and stored under vacuum at room temperature. Dimethylformamide (DMF), tetrahydrofuran (THF), and polyethyleneimine (PEI) were purchased from Sigma-Aldrich. Heparin sodium salt powder from porcine intestinal mucosa (CAS Number: 9041-08-1) was purchased from VWR (now Avantor). 1-Ethyl-3-(3-dimethylaminopropyl) carbodiimide (EDC), N-hydroxysuccinimide (NHS), and Nanopure™ water were purchased from Sigma-Aldrich. 10x Phosphate-Buffered Saline (PBS) was purchased from Thermo Fisher Scientific and diluted with deionized (DI) water.

#### S.1.2. In Vitro Biological Compatibility

250 mL prefilled Citrate Phosphate Dextrose Adenine-1 (CPDA-1) blood collection bags from Jorgensen Laboratories. Calcium Chloride Anhydrous, Reagent Grade, Innovating Science®, USA. Human Umbilical Vein Endothelial Cells (HUVECs, neonatal; Cat. No: 200-05N, Lot No: 3455) were purchased from Sigma-Aldrich and stored at –196 °C in liquid nitrogen. HUVECs at passage 4-5 were used in the experiments. The Human Endothelial Cell Growth Medium Kit (Cat. No: 211-500) was purchased from PromoCell. Trypsin-ethylenediaminetetraacetic acid (EDTA, 0.5%, no phenol red; Cat. No: 15400054), Dulbecco's Phosphate-Buffered Saline (DPBS, no calcium, no magnesium; Cat. No: 14190144, for cytotoxicity assays), and Penicillin-Streptomycin (Pen-Strep, 10 000 U/mL; Cat. No: 15-140-122) were manufactured by Gibco™ and supplied by Thermo Fisher Scientific. Gore-Tex Cardiovascular Patch (Cat. No: 1705007506) was purchased from Synergy Surgical. CyQUANT™ LDH Cytotoxicity Assay (Cat. No: C20301), CyQUANT™ Cell Proliferation Assay (Cat. No: C7026), and CyQUANT™ XTT Cell Viability Assay (Cat. No: X12223) were manufactured by Invitrogen™ and supplied by Thermo Fisher Scientific. CellCrown™ 12 well plate inserts (Cat. No: Z742383-6EA) were supplied by Sigma-Aldrich. Formalin solution (10%, neutral buffered; Cat. No: HT501128-4L), Triton X-100 (Cat. No: T9284-100ML), 4′,6-Diamidino-2-phenylindole dihydrochloride (DAPI; Cat. No: D8417-1MG), and TWEEN® 20 (suitable for cell culture; Cat. No: P2287-100ML) was supplied by Sigma-Aldrich. p120 Antibody (G-7) Alexa Fluor® 488 (Cat. No: sc-373751 AF488) was used in a 1:400 dilution

and purchased from Santa Cruz Biotechnology. SuperBlock™ Blocking Buffer (Cat. No: 37580) was purchased from Thermo Fisher Scientific.

**S.2. Supporting Figures**

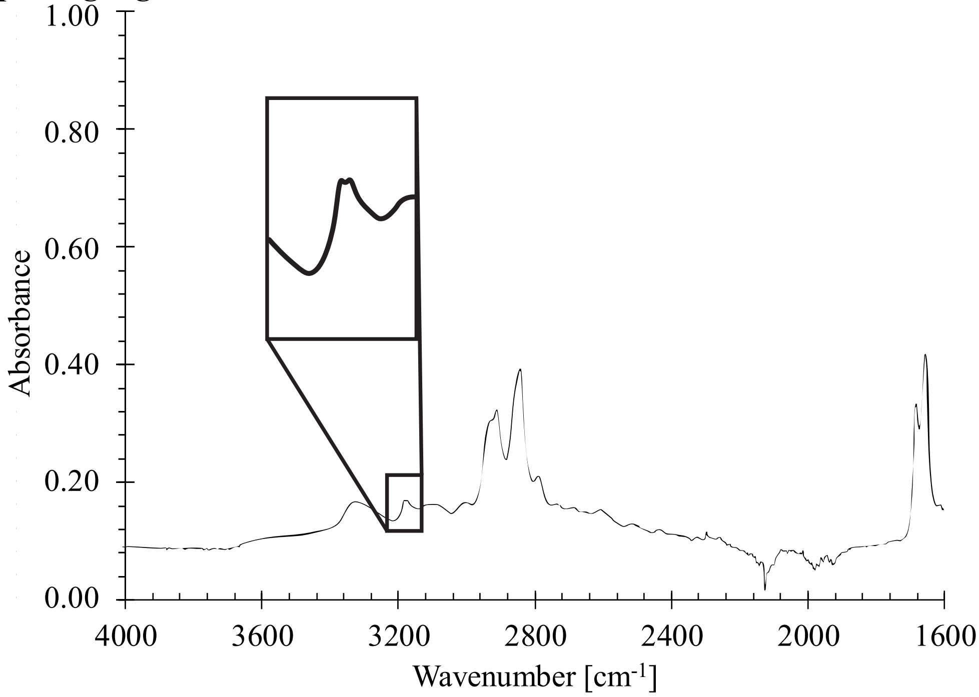


**Figure S1.** FTIR spectrum of P-ENG. The full spectrum is shown from 1600 to 4000 cm$^{-1}$, with an inset highlighting the 3100-3200 cm$^{-1}$ region. Distinct peaks in this range, corresponding to N–H and O–H stretching vibrations typical of amines and alcohols, confirm the presence of PEI in the P-ENGs.

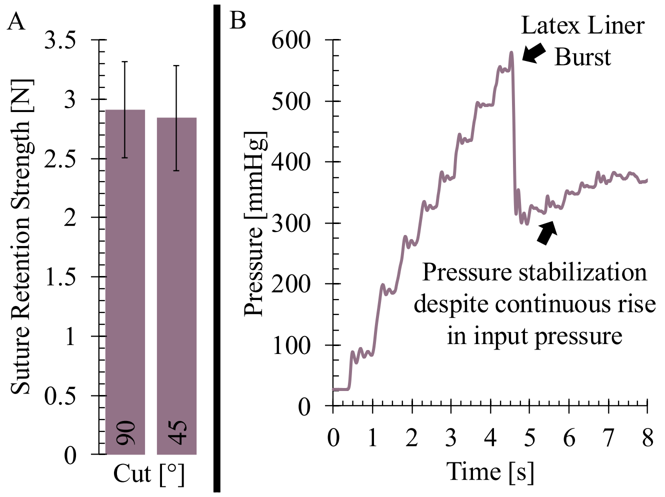


**Figure S2.** A) Evaluation of suture retention strength shows the maximum force required to pull the suture out of the material for 90° and 45° cut angles (n = 3 with 3 technical replicates each). B) Plot showing evaluation of the burst pressure shows a peak indicating where the latex liner burst occurred. Subsequent increase in input pressure indicates that due to high permeability, the ENG is unable to retain pressure above 400 mmHg (n = 3 with 3 technical replicates each).

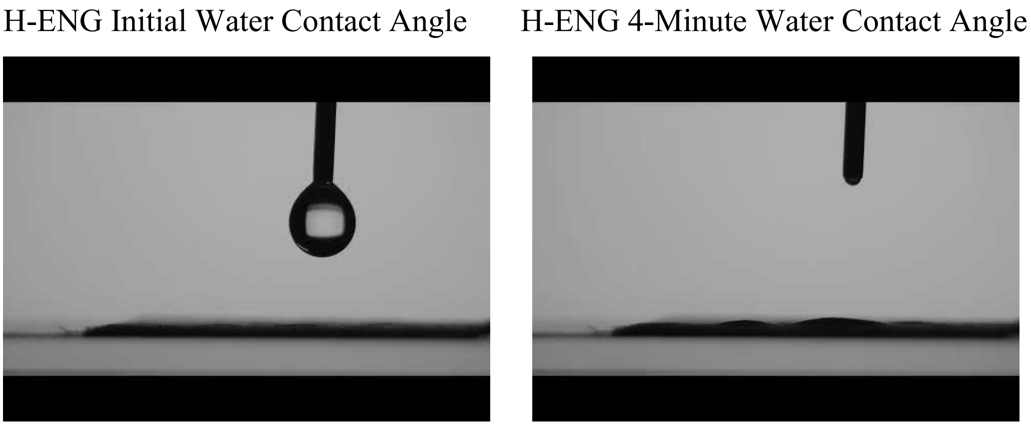


**Figure S3.** Time-dependent water contact angle of the H-ENG. Representative video of initial contact angle of 90° (left) gradually decreases to ~0° after 4 minutes (right), demonstrating complete wetting and reduced long-term hydrophobicity.

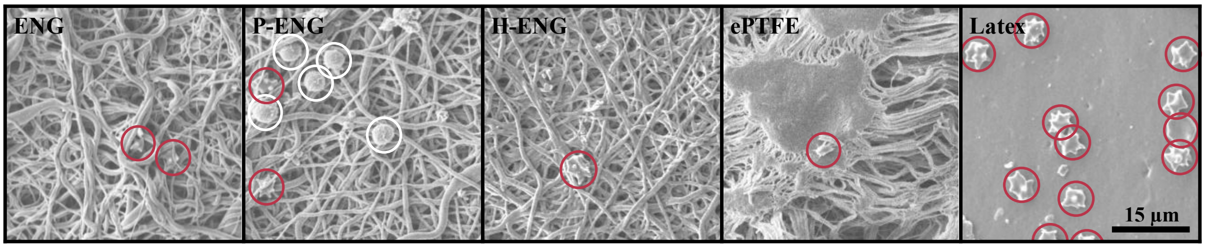


**Figure S4.** Representative SEM images of platelet adhesion assessments of ENG, P-ENG, H-ENG, ePTFE, and latex. P-ENG samples show extensive platelet (red circles) and white blood cell (white circles) adhesion, whereas H-ENG and ePTFE exhibit minimal cellular attachment.

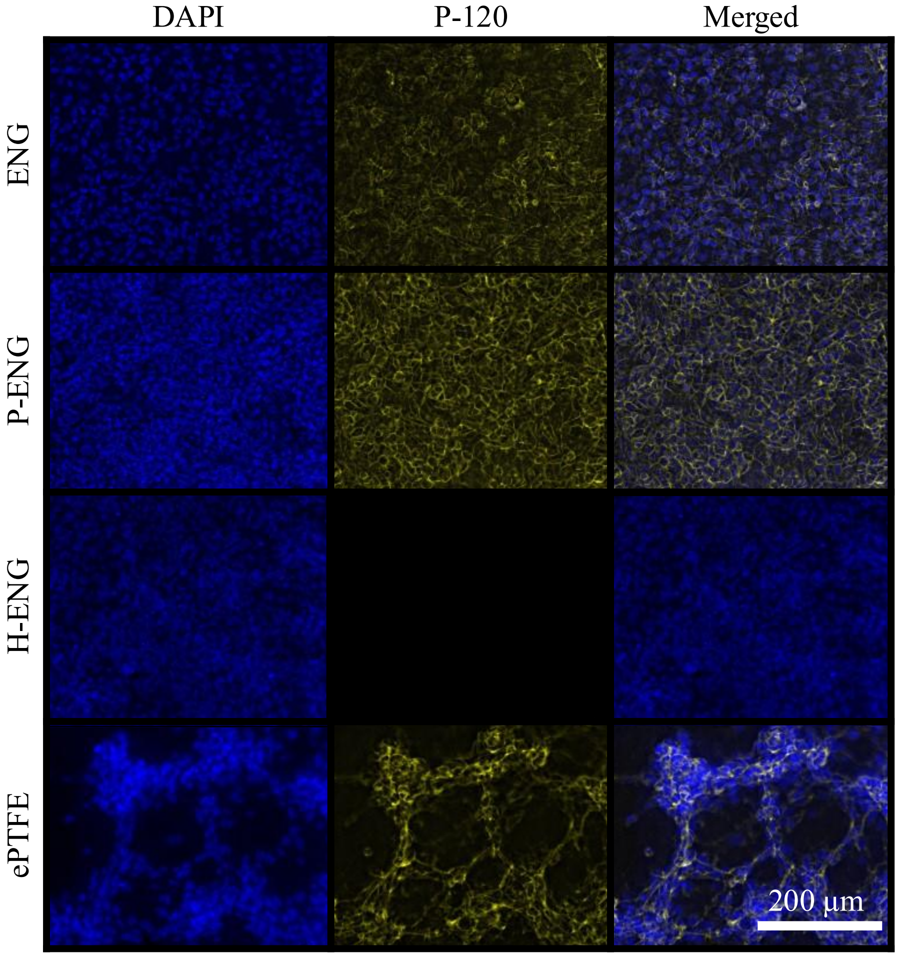


**Figure S5.** Representative fluorescence microscopy images of HUVEC cell cultures, assessing cell adhesion, morphology, and protein expression p-120 catenin (p-120) on different substrates (ENG, P-ENG, H-ENG, and ePTFE). DAPI staining highlights cell nuclei (left), while p-120 shows cell-cell junction protein distribution (middle), and the "Merged" panel overlays both signals (right). p-120 signal was not detected on H-ENG, possibly due to protein conformational masking, low expression levels, or interference from the substrate's chemical properties. *(1)*

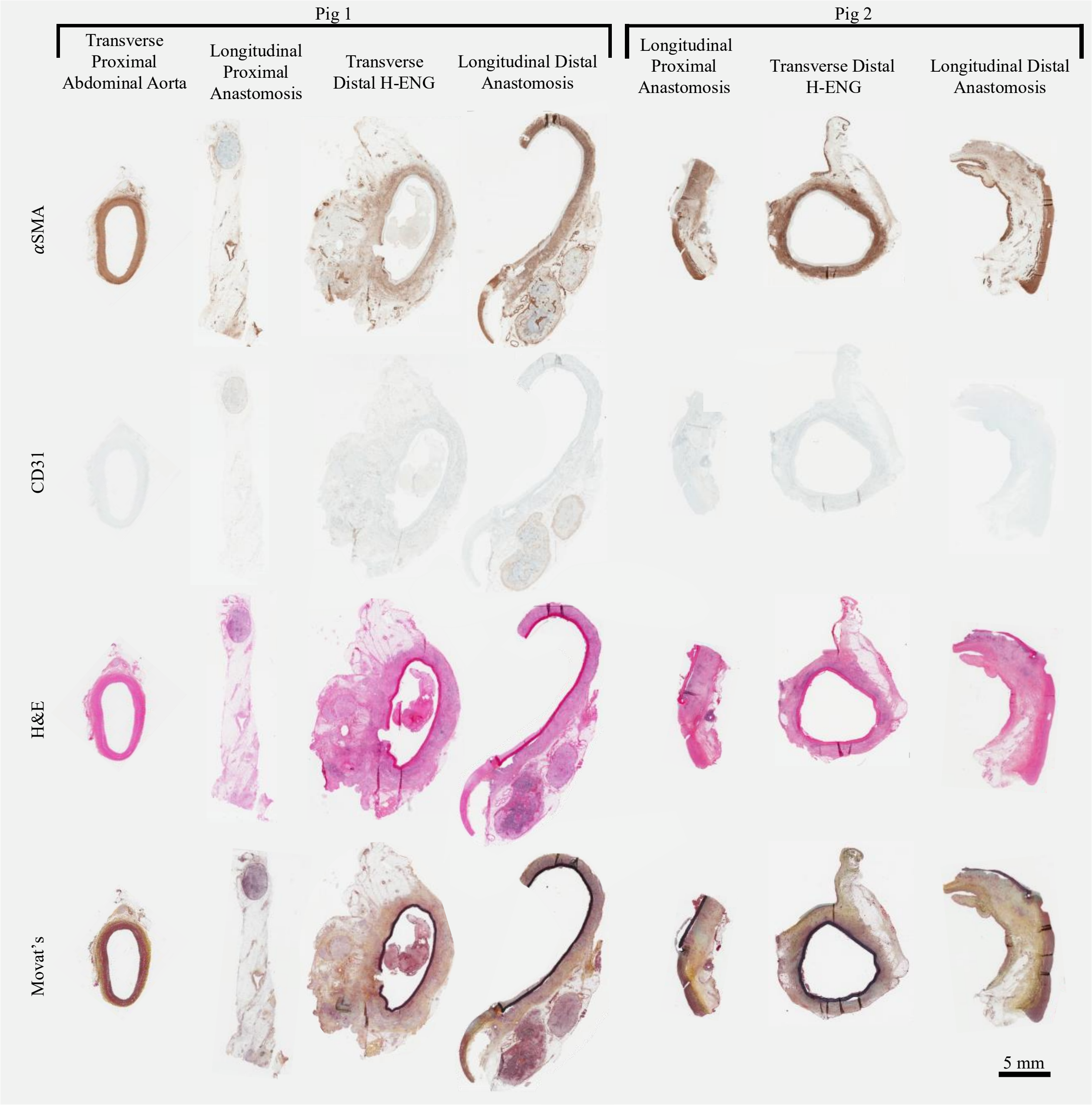


**Figure S6.** Complete histological analysis from two in vivo porcine implantation models. Rows represent different histological stains: αSMA (smooth muscle marker), CD31 (endothelial marker), H&E (general morphology), and Movat's (tissue composition). Columns correspond to anatomical locations: proximal abdominal aorta, proximal anastomosis, distal H-ENG, and distal anastomosis. For Pig 1, the proximal anastomosis section and for Pig 2, the distal anastomosis section were misaligned during tissue processing, resulting in the H-ENG graft not being visible in those samples. A localized thrombus is visible in the cross-sectional image of Pig 1's graft, positioned near the anastomosis. The remainder of the graft was free of thrombus, suggesting that the thrombus likely formed due to localized surgical manipulation rather than as a result of adverse blood-material interactions. Histology panels in their full resolution are available in figshare https://doi.org/10.6084/m9.figshare.30179797.

#### S.3. Supporting Tables

**Table S1.** Descriptive statistics (average ± standard deviation), for microstructure and mechanical properties. Average ± standard deviation for fiber diameter, graft thickness, axial stiffnesses ($K_x$), and circumferential stiffnesses ($K_y$). For all variables, n = 3 per group, where all measurements were triplicated to control for variance in electrospun fabric.

| | ENG | P-ENG | H-ENG |
|---|---|---|---|
| Fiber Diameter [µm] | 0.55 ± 0.27 | 0.61 ± 0.21 | 0.59 ± 0.26 |
| Graft Thickness [µm] | 268.89 ± 42.64 | 246.67 ± 27.13 | 265.74 ± 46.66 |
| $K_x$ [N/m] | 501.5 ± 92.9 | 480.9 ± 55.4 | 570.8 ± 54.9 |
| $K_y$ [N/m] | 311.2 ± 27.6 | 334.5 ± 36.2 | 365.3 ± 46.2 |

**Table S2.** Descriptive statistics (average ± standard deviation), for hemolysis, platelet adhesion, and static whole blood clotting assays (n = 3 with three triplicates each for all sample types).

| | ENG | P-ENG | H-ENG | ePTFE | Latex |
|---|---|---|---|---|---|
| Hemolysis [%] | 22 ± 35 | 0 ± 24 | 2 ± 23 | 4 ± 13 | |
| Platelet Adhesion [platelets/mm$^2$] | 410 ± 61 | 1,941 ± 988 | 40 ± 2 | 44 | |
| Dry Thrombus Weight [mg/cm$^2$] | 1.2 ± 0.4 | 7.0 ± 0.8 | 0.3 ± 0.6 | 0.1 ± 0.1 | 10.1 ± 0.9 |

**Table S3:** Additional descriptive statistics (given non-normal distribution: quartile one, median, and quartile three), for hemolysis, platelet adhesion, and static whole blood clotting assays (n = 3 with three triplicates each for all sample types).

| Hemolysis Assay [%] | | | | | |
|---|---|---|---|---|---|
| | ENG | P-ENG | H-ENG | ePTFE | Latex |
| Quartile One | 4 | 0 | 0 | 6 | |
| Median | 7 | 0 | 0 | 11 | |
| Quartile Three | 33 | 9 | 9 | 11 | |
| Platelet Adhesion Assay [platelets/mm$^2$] | | | | | |
| Quartile One | 380 | 1652 | 39 | | |
| Median | 410 | 1,379 | 40 | 44 | |
| Quartile Three | 441 | 2,231 | 41 | | |
| Dry Thrombus Weight [mg/cm$^2$] | | | | | |
| Quartile One | 1.0 | 6.7 | 0 | 0.1 | 9.7 |
| Median | 1.2 | 7.1 | 0 | 0.1 | 10.2 |
| Quartile Three | 1.5 | 7.5 | 0.5 | 0.2 | 10.6 |

**Table S4.** Descriptive statistics (average ± standard deviation), for all cytotoxicity assays. CyQUANT™ Cell Proliferation Assay, CyQUANT™ LDH Cytotoxicity Assay, and CyQUANT™ XTT Cell Viability Assay results of HUVECs cultured on ENG-type materials, ePTFE negative control (NC), and positive control (PC). n = 3 per group, where all assays were repeated four times per sample.

| CyQUANT™ XTT Cell Viability Assay: Absorbance [450] | | | | | | |
|---|---|---|---|---|---|---|
| | ENG | P-ENG | H-ENG | ePTFE | NC | PC |
| Mean | 1.257 | 1.009 | 0.888 | 0.896 | 0.792 | 0.092 |
| Standard Deviation | 0.310 | 0.184 | 0.124 | 0.101 | 0.063 | 0.001 |
| CyQUANT™ LDH Cytotoxicity Assay: Absorbance [490/680] | | | | | | |
| Mean | 0.033 | 0.035 | 0.030 | 0.034 | 0.034 | 0.248 |
| Standard Deviation | 0.002 | 0.003 | 0.002 | 0.002 | 0.002 | 0.183 |
| CyQUANT™ Cell Proliferation Assay: Viability [%] | | | | | | |
| Mean | 96.049 | 92.816 | 95.594 | 86.207 | 100.000 | 10.129 |
| Standard Deviation | 7.011 | 7.130 | 5.844 | 10.836 | 14.815 | 2.760 |

**Table S5.** Duplex ultrasound evaluation of H-ENG and abdominal aorta. Representative transverse and longitudinal duplex ultrasound videos showing the H-ENG and the abdominal aorta in Pig 1 and 2 at two time points: implantation and two weeks post-implantation. The transverse view compares the native abdominal aorta (left) and the H-ENG. The longitudinal view shows proximal anastomosis and distal anastomosis (right) of the H-ENG (right in proximal anastomoses ultrasound and left in distal anastomoses). The two-week distal anastomosis ultrasound of Pig 1 shows a localized thrombus with oscillatory movement within the lumen. All ultrasound assessments confirm that H-ENGs maintain pulsatile behavior during the cardiac cycle.

| | | Transverse View | | Longitudinal View | |
|---|---|---|---|---|---|
| | | Abdominal Aorta | H-ENG | Proximal Anastomosis | Distal Anastomosis |
| Pig 1 | Implantation | 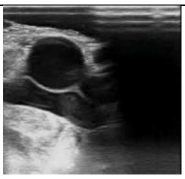 | 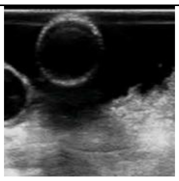 | 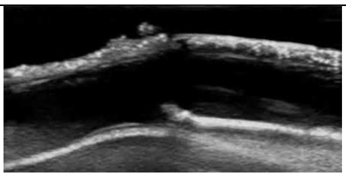 | 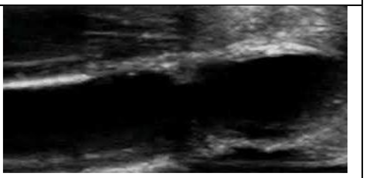 |
| | Two-week | 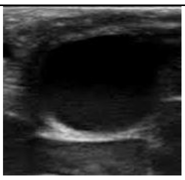 | 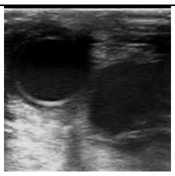 | 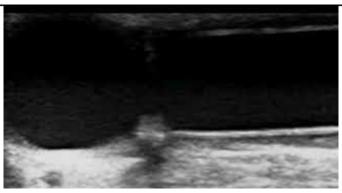 | 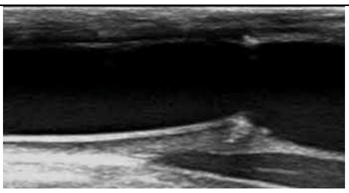 |
| Pig 2 | Implantation | 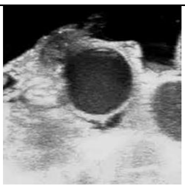 | 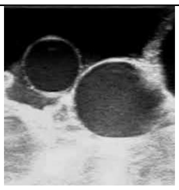 | 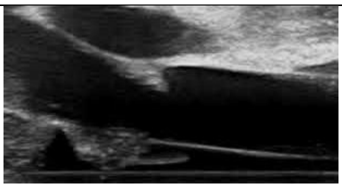 | 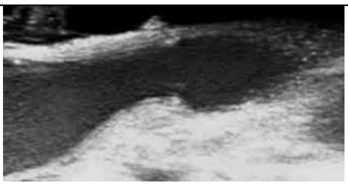 |
| | Two-week | 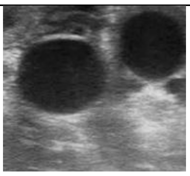 | 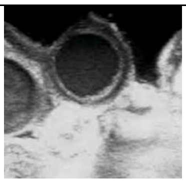 | 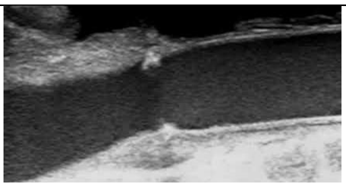 | 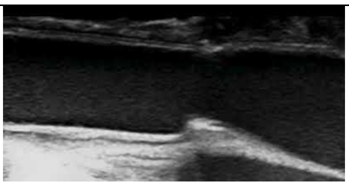 |

**Table S6.** Duplex ultrasound with flow of H-ENG and abdominal aorta. Representative longitudinal duplex ultrasound videos of the H-ENG in Pig 1 and Pig 2 at two time points: implantation and two weeks post-implantation. The longitudinal view shows the proximal anastomosis (left) and distal anastomosis (right) of the H-ENG. For Pig 2, the two-week distal anastomosis image was not available. All ultrasound assessments demonstrate maintained flow patterns at both anastomotic sites and confirm that H-ENGs preserve pulsatile behavior during the cardiac cycle.

<table>
<tr><th colspan="2" rowspan="2"></th><th colspan="2">Longitudinal View</th></tr>
<tr><th>Proximal Anastomosis</th><th>Distal Anastomosis</th></tr>
<tr><td rowspan="2">Pig 1</td><td>Implantation</td><td>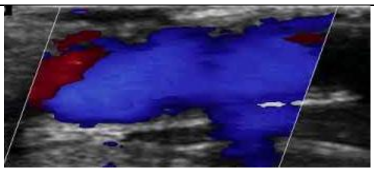</td><td>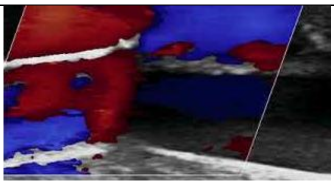</td></tr>
<tr><td>Two-week</td><td>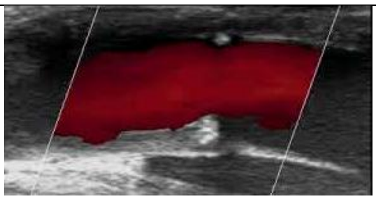</td><td>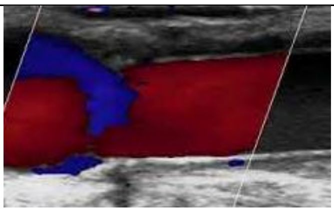</td></tr>
<tr><td rowspan="2">Pig 2</td><td>Implantation</td><td>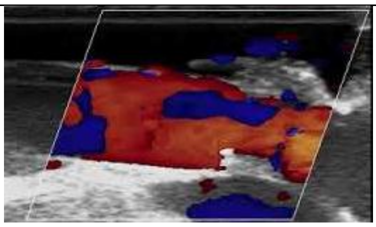</td><td>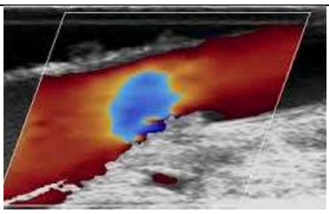</td></tr>
<tr><td>Two-week</td><td>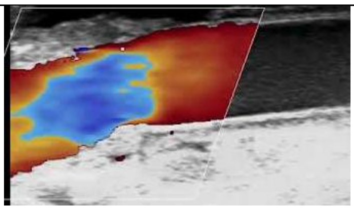</td><td>Not available.</td></tr>
</table>

**References**

1. Piña, R.; Santos-Díaz, A. I.; Orta-Salazar, E.; Aguilar-Vazquez, A. R.; Mantellero, C. A.; Acosta-Galeana, I.; et al. Ten Approaches That Improve Immunostaining: A Review of the Latest Advances for the Optimization of Immunofluorescence. *International Journal of Molecular Sciences* **2022**, *23*(3), 1426. doi:10.3390/ijms23031426.
